# Lifetime fitness benefits of short-distance dispersal are associated with inbreeding tolerance despite multiple inbreeding avoidance mechanisms

**DOI:** 10.1101/2024.09.21.614049

**Authors:** Shailee S. Shah, Jennifer Diamond, Sahas Barve, Elissa J. Cosgrove, Reed Bowman, John W. Fitzpatrick, Nancy Chen

## Abstract

It is commonly assumed that mechanisms that reduce inbreeding, such as sex-biased dispersal or kin avoidance, are always adaptive. However, theoretical models predict that inbreeding avoidance evolves only when the relative costs of inbreeding are higher than the costs of inbreeding avoidance. We empirically tested this prediction with 32 years of data on Florida scrub-jays (*Aphelocoma coerulescens*), birds with short dispersal distances and multiple mating opportunities over their lifetimes. Using simulations, we showed that limited dispersal increases the inbreeding risk by at least two-fold, with a stronger effect at later life stages. Sex-biased dispersal reduces, but does not eliminate, this risk. Additionally, we found that the fitness costs of inbreeding are lower than those of dispersal in terms of lifetime reproductive success (LRS) and survival. Inbreeding is associated with substantially lower LRS only when mates are first-order kin. Accordingly, we found that the population shows a degree of inbreeding tolerance but, within their dispersal constraints, Florida scrub-jays actively avoid mating with first-order kin. Overall, our results empirically demonstrate the context-specific nature of inbreeding avoidance, with selection favoring limited dispersal despite costs of inbreeding and inbreeding avoidance via mate choice only when the relative fitness costs of inbreeding are particularly high.

## Introduction

Though it is commonly assumed that organisms should avoid mating with relatives, or inbreeding, which can have negative fitness consequences (Pusey and Wolf 1996), theoretical models repeatedly suggest that inbreeding avoidance should be highly context dependent (Bengtsson 1978; Duthie and Reid 2016; Kokko and Ots 2006; Waser et al. 1986). If inbreeding avoidance has associated fitness costs—such as loss of breeding opportunities, movement to suboptimal habitat, heightened mortality risk, or outbreeding depression (Griesser et al. 2006; Hu et al. 2017; Jain 1976; O’Riain et al. 2000; Peer and Taborsky 2005)— that are higher than the fitness costs of inbreeding, certain levels of inbreeding may instead be tolerated (Kokko and Ots 2006; Waser et al. 1986) or even favored for its indirect fitness benefits (Gow et al. 2019). Indeed, studies across a broad range of taxa show little evidence of inbreeding avoidance (de Boer et al. 2021; Goodwillie et al. 2005). However, while the conditions where inbreeding is tolerated or preferred (*i.e.*, inbreeding occurs as expected by chance or more than expected by chance) are well understood in other taxa, such as selfing in plants (Jarne and Charlesworth 1993; Karron et al. 2012) and fungi (Nieuwenhuis and James 2016), studies in animals are still heavily biased towards the assumption that any degree of inbreeding is maladaptive (de Boer et al. 2021). To challenge this assumption, we need a more nuanced investigation of life history traits that determine conditions favoring evolution of inbreeding avoidance strategies and the fitness outcomes that determine their persistence (de Boer et al. 2021; Kokko and Ots 2006).

Dispersal and mate choice are fundamental life history traits that govern the likelihood of encountering and choosing relatives as mates, thereby influencing the scope for inbreeding avoidance (Blouin and Blouin 1988; Perrin and Mazalov 1999). While dispersal can effectively eradicate inbreeding risk if all individuals move far enough away to eliminate any likelihood of encountering relatives as potential mates (Gandon 1999), dispersal is often costly when subject to constraints such as finite time and energy, increased mortality risk, or opportunity costs (Ashman and Schoen 1996; Bonte et al. 2012).

Considerable variation exists in the propensity and distance of dispersal both within and across species (Comte and Olden 2018; LaRue et al. 2018; Schloss et al. 2012; Stevens et al. 2010). In cases where the propensity for and/or distance of dispersal is low—*i.e.*, dispersal is “limited” (Shields 1982)—variation in dispersal between the sexes (*i.e.*, sex-biased dispersal) mostly mitigates inbreeding risk (Li and Kokko 2019; Perrin and Mazalov 1999). However, some species face constraints on sex-biased dispersal such as range-restriction (Alzate and Onstein 2022), low propagule dispersal potential (Lloyd et al. 2018), or social structure (Nichols 2017). In such species, the scope for the evolution of active inbreeding avoidance mechanisms via mate choice, such as self-incompatibilities (Takayama and Isogai 2005), multiple mating or mate switching (Cockburn et al. 2003; Tregenza and Wedell 2002), and kin avoidance (Komdeur and Hatchwell 1999; Leedale et al. 2024) may be higher.

The scope for inbreeding avoidance may also vary at different life stages in species that experience multiple mate choice events (Szulkin et al. 2013). The benefits of mate choice can vary with age and experience such that individuals become more or less choosy at later life stages (Edward and Chapman 2013), with sequential matings (Gabor and Halliday 1997; Pitcher et al. 2003), or when switching to a different pair-bonded mate (Hidalgo Aranzamendi et al. 2016; Otter and Ratcliffe 1996). While inbreeding avoidance has not been specifically examined in this context, its relative fitness outcomes may also shift with life stages. For example, in a territorial species, moving away from an established breeding territory for any subsequent pairings may carry an increased mortality risk (Daniels and Walters 2000a) and loss of fitness benefits such as site familiarity and reduced competitive interactions (Beck et al. 2020; Byrne et al. 2022; Piper et al. 2023). Though limited dispersal away from the breeding territory—*i.e.*, “site fidelity” (Switzer 1993)—also can increase inbreeding risk when local relatedness increases with time (Croft et al. 2021), inbreeding tolerance may be higher at later life stages if fitness benefits outweigh costs of inbreeding. Understanding when and why inbreeding avoidance strategies evolve thus requires examining dispersal and mate choice across individuals’ lifetimes, not just at the natal dispersal (*i.e.*, movement for first pairing) and first mate choice stages.

For long-lived species that may mate with multiple individuals in their lifetimes, understanding the conditions that foster the evolution of inbreeding avoidance involves five key components: (i) sufficient variation in relatedness between potential mates; (ii) researchers’ ability to identify actual and (iii) potential mates; (iv) accurate measures of relatedness; and (v) full accounting of mate choice events occurring throughout different stages of an individual’s lifetime (Szulkin et al. 2013). Though previous studies of inbreeding have incorporated one or several of these components, to our knowledge no single study meets all five requirements (Gibbs and Grant 1989; Wheelwright and Mauck 1998; Daniels and Walters 2000b; Foerster et al. 2006; Frommen and Bakker 2006; Szulkin et al. 2009; Billing et al. 2012; Galezo et al. 2022). Additionally, to understand why a certain degree of inbreeding avoidance does or does not exist, measures of lifetime fitness are required. Variable reproductive strategies over individuals’ lifetimes may compensate for any short-term fitness costs associated with inbreeding or inbreeding avoidance strategies, especially in long-lived species (Van de Casteele et al. 2003).

Here, we leveraged 32 years of individual-level data on dispersal, mating, survival, and reproductive success from a long-lived avian species, the Florida scrub-jay (*Aphelocoma coerulescens*), to investigate how dispersal and mate choice decisions contribute to inbreeding risk and the conditions underlying the evolution of inbreeding avoidance in a natural population. Breeding pairs of Florida scrub-jays defend year-round territories, and most offspring delay breeding for one or more years, partly as a mate choice tactic (Woolfenden and Fitzpatrick 1984). Offspring typically disperse less than 2.5 km away (or the equivalent of one to five territories), and occasional long-distance dispersal events suggest that this pattern of limited dispersal is not due to anatomical restrictions (Suh et al. 2020). Dispersal is sex-biased, with females dispersing farther than males (Suh et al. 2020; Woolfenden and Fitzpatrick 1984). Relatedness of potential mates within a population ranges from parents/full-siblings to unrelated immigrants (Aguillon et al. 2017). Evidence of inbreeding depression suggests that inbreeding avoidance strategies may be adaptive in this species (Chen et al. 2016), though long-distance dispersal may also have associated fitness costs (Fitzpatrick et al. 1999).

We sought to test the theoretical prediction that evolution of inbreeding avoidance is context dependent and not always adaptive (Bengtsson 1978; Duthie and Reid 2016; Kokko and Ots 2006; Waser et al. 1986). Our specific objectives were to investigate the influence of dispersal on the scope for the evolution of inbreeding avoidance, relative fitness consequences of inbreeding versus inbreeding avoidance that may contribute to its persistence, and degree of inbreeding avoidance via mate choice. First, to examine the extent to which limited dispersal increases inbreeding risk, we quantified the expected relatedness of breeders of both sexes to mates under three random mating scenarios with increasing constraints on dispersal: (i) no constraints on dispersal distance, (ii) limited dispersal distance with existing sex bias included, and (iii) limited dispersal distance with any sex bias removed. With this study design, we were able to examine how both no constraints on dispersal distance for either sex (model i) and equal dispersal constraints on both sexes (model iii) changed the inbreeding risk compared to the observed pattern of sex-biased dispersal constraints (model ii). We predicted that expected relatedness to mates would be higher with increasing dispersal constraints for both sexes across all life stages (*i.e.*, model iii > model ii > model i). Further, we predicted that if the observed pattern of sex-biased dispersal mitigates inbreeding risk, simulated relatedness to mates will be lower in scenario ii than scenario iii. In fact, if sex-biased dispersal is sufficient to negate inbreeding risk, simulated relatedness to mates would be roughly equivalent between scenarios i and ii.

Next, we quantified fitness consequences of inbreeding versus inbreeding avoidance via dispersal. We predicted that both breeding with related individuals and dispersing long distances would lead to lower lifetime fitness for both sexes at both life stages. Finally, based on the relative fitness outcomes, we predicted that if the cost of inbreeding is greater than or equivalent to the cost of long-distance dispersal, active inbreeding avoidance will exist. If the fitness costs of inbreeding are severe even when mates are distantly related, we expected to find strict kin avoidance. In this case, observed relatedness to mates should approach the level expected under random mating with unconstrained dispersal (*i.e.*, scenario i). Alternatively, if the cost of breeding with moderately related individuals is low, we expected to find a degree of inbreeding tolerance. In this case, observed relatedness to mates will be higher than expected under scenario i but lower than expected under random mating within the observed dispersal patterns (*i.e.*, scenario ii). Further, if kin avoidance is facilitated by kin recognition, the observed proportion of mates in categories of close kin (first-, second- and third-order kin) should follow the same patterns. We also examined mate switching and predicted that pairs with more closely related mates will be more likely to dissolve because of mate switching than because of death of a mate.

## Methods

### Data collection

#### Study site and species

Florida scrub-jays at Archbold Biological Station (27.10°N, 81.21°W) have been continuously monitored since 1969, with the study tract expanded to its current size (mean ± sd = 914.04 ± 143.44 ha) by 1990 (Woolfenden and Fitzpatrick 1984). Detailed monitoring data from this population provides extraordinary ability to identify all actual and potential mates, calculate accurate relatedness measures using a single, extensive pedigree, incorporate mate choice events throughout individuals’ lifetimes, and quantify lifetime fitness in terms of reproductive success and survival (Fitzpatrick and Bowman 2016). All birds are banded with unique combinations of color bands. Throughout the breeding season (early March to early June), all nests are located and monitored for fledging or failure. All territories are mapped in April, with territory boundaries documented at locations of territorial disputes between neighboring groups. In addition to nest monitoring, the population is censused each month, resulting in detailed data on dispersal and survival of both fledglings and adults. Pairs are socially monogamous, with exceptionally low rates of extra-pair paternity (<1%) (Townsend et al. 2011), allowing accurate parentage inference from field observations alone. Furthermore, previous work verified genetic parentage of offspring born between ∼1989-1991, 1994-1995, and 1999-2013 (Chen et al. 2016). Using these detailed data, we have accumulated a 15-generation pedigree of known-age, known-parentage individuals with precise measurements of lifespan and reproductive success. For this study, we used data from 1990-2021, which included on average (± SD) 71.4 (± 14.7) territories (area mean ± sd =13.24 ± 6.37 ha) per year. Fieldwork was approved by the Institutional Animal Care and Use Committees at Cornell University (IACUC 2010-0015) and Archbold Biological Station (AUP-006-R) under permits from the US Fish and Wildlife Service (TE824723-8, TE-117769), the US Geological Survey (banding permits 07732, 23098), and the Florida Fish and Wildlife Conservation Commission (LSSC-10-00205).

#### Observed mates

Our dataset of observed mates included all new breeding pairs formed between 1990 and 2021 (*N* = 1,077). New breeding pairs include pairs formed by two first-time breeders, one first-time and one experienced breeder, or two experienced breeders. For first-time breeders, we called the event a “first pairing” and for experienced breeders, a “later pairing”. The first pairing for one mate is thus not always the first pairing for the other mate (*e.g.*, an experienced male could re-pair with a female who is a first-time breeder, and *vice versa*). New pairs might occupy a new territory, which may be inherited by one of the two breeders or formed as an extension of their natal territory in a process called “budding” (Woolfenden and Fitzpatrick 1984; Fitzpatrick and Bowman 2016; Beres et al. 2026). Alternatively, a new pair might form on a non-natal territory already occupied by one of the two breeders, where a breeding vacancy is typically formed due to the death of a breeder, but occasionally via mate switching (Woolfenden and Fitzpatrick 1984; Fitzpatrick and Bowman 2016). Thus, both first and later pairings can occur without any movement away from the currently occupied territory by at least one of the two breeders.

#### Observed dispersal distance

We measured the distance moved for pairing as the number of territories crossed, which is the most biologically relevant measure of distance moved for a territorial species (Fitzpatrick et al. 1999; Greenwood and Harvey 1982). The number of territories crossed was highly correlated with distance moved in meters (Pearson’s *r* = 0.91; Fitzpatrick et al. 1999). Henceforward, we refer to “number of territories crossed” as “dispersal distance” for brevity. We drew a straight-line movement trajectory from the centroid of an individual’s previous territory to the centroid of its new breeding territory. We then used the *st_intersects* function from the R package *sf* (Pebesma 2018) to calculate the number of territories that each line intersected (see *SI* for details). When the focal individual remained on its natal territory and became a breeder, formed a territory by budding, or remained on its non-natal breeding territory, we quantified distance moved as 0 territories crossed. To obtain accurate measures of distance moved, we excluded any pairings that involved movement that started or ended outside of the study tract.

#### Observed relatedness to mates

To quantify the observed level of inbreeding, we calculated the kinship coefficient between mates using the population pedigree with the kinship function in the R package *kinship2* (Sinnwell et al. 2014). With no inbreeding, the kinship coefficient ranges from 0 to 0.5, such that 0.5 is an individual’s kinship coefficient with itself; 0.25 is the average kinship coefficient between siblings and between parents and offspring (first-order kin); 0.125 is the average kinship coefficient between half-siblings, grandparents-grandchildren, and aunt/uncle-niece/nephew pairs (second-order kin); and 0.0625 is the average kinship coefficient between first cousins (third-order kin) (Malécot 1948). Pedigree-based inbreeding estimates are correlated with genomic-based estimates in our study population, confirming that our pedigree is complete enough to provide accurate measures of inbreeding and relatedness (Chen et al. 2016).

### Data analysis

All analyses were conducted using R version 4.2.2 (R Core Team 2022). Fixed effects for all models were standardized using z scores (Schielzeth 2010), and the variance inflation factor was < 2 for all fixed effects in all GLMMs (Fox and Monette 1992).

### Influence of dispersal on the scope for the evolution of inbreeding avoidance

To simulate and quantify the effect of limited dispersal on inbreeding risk, we constructed three random mating scenarios with (i) no constraints on dispersal distance, (ii) limited dispersal distance with existing sex bias included, and (iii) limited dispersal distance without sex bias (Figs. S1-S2). For the unconstrained model, we defined potential mates for all individuals of one sex in a given year as all individuals of the opposite sex from our dataset of observed mates (*i.e.*, individuals that were both available as potential mates and successful at finding new mates) (*N* focal individuals: mean ± SD = 25.4 ± 7.06; *N* available mates per individual: mean ± SD = 25.5 ± 7.01; see *SI* for details). For the limited dispersal with sex-biased dispersal model, we limited available mates to opposite sex individuals at the observed dispersal distance of each individual (*N* available mates per individual: mean ± SD = 1.37 ± 0.63). Finally, for the limited dispersal but no sex bias model, we randomly shuffled observed dispersal distance values across sexes, but within life stages (*i.e.*, first or later pairing). We then limited available mates for each focal individual to opposite sex individuals at the assigned dispersal distance (*N* available mates per individual: mean ± SD = 1.39 ± 0.68).

For each focal individual we then randomly assigned an available mate for each year. These random assignments were permuted 1000 times. In dispersal-constrained models, individuals that did not move away from their territory but paired with a new mate were randomly assigned a simulated mate from a territory equidistant to the dispersal distance of their new mate. For example, if a male retained his territory (territory “XYZ”) but with a new female that moved across two territories to reach XYZ, we assigned the male a dispersal distance value of 0 territories crossed, and a simulated mate randomly picked from females on territories two territories away from XYZ. While both mates may move territories to form a new breeding pair (107 out of 1077 [10%] observations), our simulations are sex-specific and only account for dispersal by the focal sex.

To quantify expected levels of inbreeding under the different scenarios we calculated the kinship coefficient between simulated breeding pairs using the same method as described above. We then compared the simulated mean kinship coefficients between the three models using *t*-tests, with Bonferroni corrections to *p*-values to account for multiple testing (*N* = 1000 iterations per model). All analyses were run separately by sex and life stage.

### Fitness consequences of inbreeding versus inbreeding avoidance

To further investigate why inbreeding avoidance or tolerance may evolve and persist in our study population, we quantified the relative fitness costs associated with inbreeding and inbreeding avoidance. For these analyses, we restricted our dataset to individuals from cohorts from which all members born within the study tract were last observed before 2021. Without this restriction, our dataset would have consisted of individuals that had died (*i.e.*, short-lived individuals), but not individuals that were still living (*i.e.,* long-lived individuals) from certain birth cohorts, biasing our dataset towards short-lived individuals (Barve et al. 2021; Shah and Rubenstein 2022). The final dataset for all fitness analyses included individuals born between 1988-2006 and in 2009 (*N*: Males = 197, Females = 120).

#### Breeder survival

We assessed breeder survival using monthly census and capture records and calculated breeding lifespan as the number of years from becoming a breeder to death. To examine the effect of distance moved on breeder survival, we fit stratified Cox proportional hazards models with two time intervals: ≤ 4 years and > 4 years from the start of breeding (Zhang et al. 2018). These models accounted for a time-varying effect of distance moved such that the proportional hazard assumption was met (see *SI* for details, Table S1, Fig. S3). We used the *Surv* and *coxph* functions from the R package *survival* (Therneau 2022) to fit separate models by sex of breeder, and incorporated natal year as a random effect (*N*: Males = 197, Females = 120). For the subset of individuals who switched mates at least once in their lifetime (*N*: Males = 91, Females = 50), we also examined the effect of site fidelity on breeder survival using Cox proportional hazards models with site fidelity (1 = site unfaithful, 0 = site faithful) as a fixed effect and natal year as a random effect.

#### Lifetime reproductive success

We calculated lifetime reproductive success (LRS) for all breeders as the total number of offspring produced over an individual’s lifetime that survived to adulthood, *i.e.*, age 1 year, the stage at which Florida scrub-jays become physiologically capable of breeding (Schoech et al. 1996). Since the mean growth rate of our study population is close to 1 (0.998) (Summers et al. 2024), we deemed LRS an appropriate metric of individual fitness (Brommer et al. 2002). To examine how distance moved for pairing and inbreeding impact lifetime fitness of Florida scrub-jays, we fit two separate GLMMs by sex with negative binomial error distributions (to account for overdispersion), LRS as the dependent variable, and distance moved for first pairing, weighted relatedness to mates, and breeding lifespan as fixed effects (*N*: Males = 197, Females = 120). We calculated a weighted kinship coefficient to mates for all breeders across their lifetimes by multiplying the kinship coefficient of the focal individual to each of their mates by the number of years they were paired with that mate and summing the resulting values (Fig. S4). Breeding lifespan was included in the model to account for the fact that the more breeding seasons a breeder experiences, the more chances it has at successful reproduction (Fitzpatrick and Bowman 2016). We also included natal year as a random effect to account for variation between birth cohorts. We used the function *glmer.nb* in the R package MASS (Venables and Ripley 2002).

Additionally, to specifically examine the effect of moving away from the breeding territory, we also fit two additional GLMMs with the same dependent variable and total distance moved for later pairings, weighted kinship coefficient to mates, and breeding lifespan as fixed effects using a truncated dataset that only included individuals that had switched mates in their lifetimes (*N*: Males = 91, Females = 50). For females, we used a zero-inflated Poisson error distribution since the model did not converge with a negative binomial error distribution due to the small sample size. We included only breeding lifespan, which has the strongest effect on the proportion of individuals with LRS = 0, as a fixed effect for the zero-inflation component (adding additional fixed effects in the zero-inflation component led to convergence issues). We used the *glmmTMB* function in the R package glmmTMB (Brooks et al. 2017). For males, we retained a negative binomial error distribution since there were no model convergence issues and the negative binomial model provided a more parsimonious fit (ΔBIC = 4.14). However, using a zero-inflated Poisson error distribution for the model for males did not change the results (Table S2). Model fits were checked using quantile-quantile plots of residuals (Fig. S5) generated using the R package DHARMa (Hartig 2026). Fixed effects were not correlated (Table S3).

### The degree of inbreeding avoidance via mate choice

To investigate the degree to which Florida scrub-jays actively avoid inbreeding via kin avoidance, we compared the observed population means of inbreeding (measured as kinship coefficient to mates), aggregated across years, to the expected distributions generated under random mating scenarios i (unconstrained dispersal) and ii (limited, sex-biased dispersal) using empirical *p*-values (Davison and Hinkley 1997). Specifically, we quantified the proportion of simulations that had a mean value less than or equal (*S*_low_) or greater than or equal (*S*_high_) to the observed mean. Empirical *p*-values were then calculated as (*S*_high/low_+1)/(*N*+1), where *N* was the total number of simulations (*i.e.,* 1000). We also compared the observed proportions of first-, second- and third-order kin mates to the expected proportions under both random mating scenarios. All analyses were run separately by sex and life stage.

To investigate whether Florida scrub-jays avoided the impact of inbreeding on reproductive success via mate switching, we fit generalized linear mixed models (GLMMs) with the cause of pair dissolution (1 = mate switching, 0 = mate death) as the dependent variable, the kinship coefficient to mate as a fixed effect, year as a random effect, and a binomial error structure. Individual ID was also included as a random effect in models of later pairings since some breeders engaged in multiple later pairings over their lifetimes. We also fit models with a categorical fixed effect of kinship between mates (1 = first-order relative, 0 = not a first-order relative) to account for cases where opposite-sex offspring might inherit dominant breeding positions, resulting in mother-son or father-daughter pairings that may be more likely to end in mate switching (Cockburn et al. 2003). We fit separate GLMMs by sex and life stage (*N*: Males, First pairing = 194; Males, Later pairing = 187; Females, First pairing = 163; Females, Later pairing = 169 pairings) using the function *glmer* in the R package *lme4* (Bates et al. 2015).

## Results

### Observed dispersal and mate choice patterns

The annual number of new breeding pairs ranged from 17-47 pairs with an average (± SD) of 29.2 (± 8.03) new pairs per year. Of the individuals included in our lifetime fitness analysis, 42% of females (50 out of 120) and 46% of males (91 out of 197) re-paired at least once in their lifetime. Only about half of all later pairings were a result of subordinates moving in to fill a breeding vacancy (see *SI* for details), though most new mates came from within a two-territory radius if the focal breeder retained its breeding territory (Fig. S6). Females, on average, moved farther than males, though the effect size was greater for first pairings than later pairings (see *SI*, Fig. S1). Across all years, only 43 out of 1077 (4%) of observed new breeding pairs consisted of individuals who were at least as closely related as first cousins (*i.e.*, had a kinship coefficient ≥ 0.0625) and only 13 out of 1077 (1.2%) of observed breeding pairs comprised first-order relatives (7 mother-son, 3 father-daughter, and 3 full sibling pairs; Figs. S2, S7). Relatedness to potential mates was significantly higher at closer geographical distances (see *SI*, Fig. S8).

### Influence of dispersal on the scope for the evolution of inbreeding avoidance

We confirmed that limited dispersal increased inbreeding risk in Florida scrub-jays. Under a model of random mating with limited but not sex-biased dispersal (scenario iii), mean relatedness to mates was two-fold or more higher for both sexes for all life stages than expected under a model of random mating with unconstrained dispersal (scenario i) (Males, First Pairings: *t*_637757_ = 68.12, *P* < 0.001; Females, First pairings: *t*_565627_ = 72.94, *P* < 0.001; Males, Later pairings: *t*_472441_ = 82.40, *P* < 0.001, Females, Later pairings: *t*_503923_ = 77.13, *P* < 0.001; Fig. 1A-D). The observed sex bias in dispersal does help mitigate inbreeding risk: the expected level of inbreeding for first pairings across both sexes was significantly lower (by ≥ 50%) under a model of random mating with limited, sex-biased dispersal (scenario ii) compared to a random mating model with limited dispersal without sex bias (scenario iii) (Males: *t*_527991_ = -44.39, *P* < 0.001; Females: *t*_520491_ = -46.64, *P* < 0.001, Fig. 1A-B) and for females for later pairings (*t*_597723_ = -13.39, *P* < 0.001, Fig.1C), but the pattern was reversed for males for later pairings (*t*_585009_ = 34.9, *P* < 0.001, Fig. 1D).

**Figure 1.**
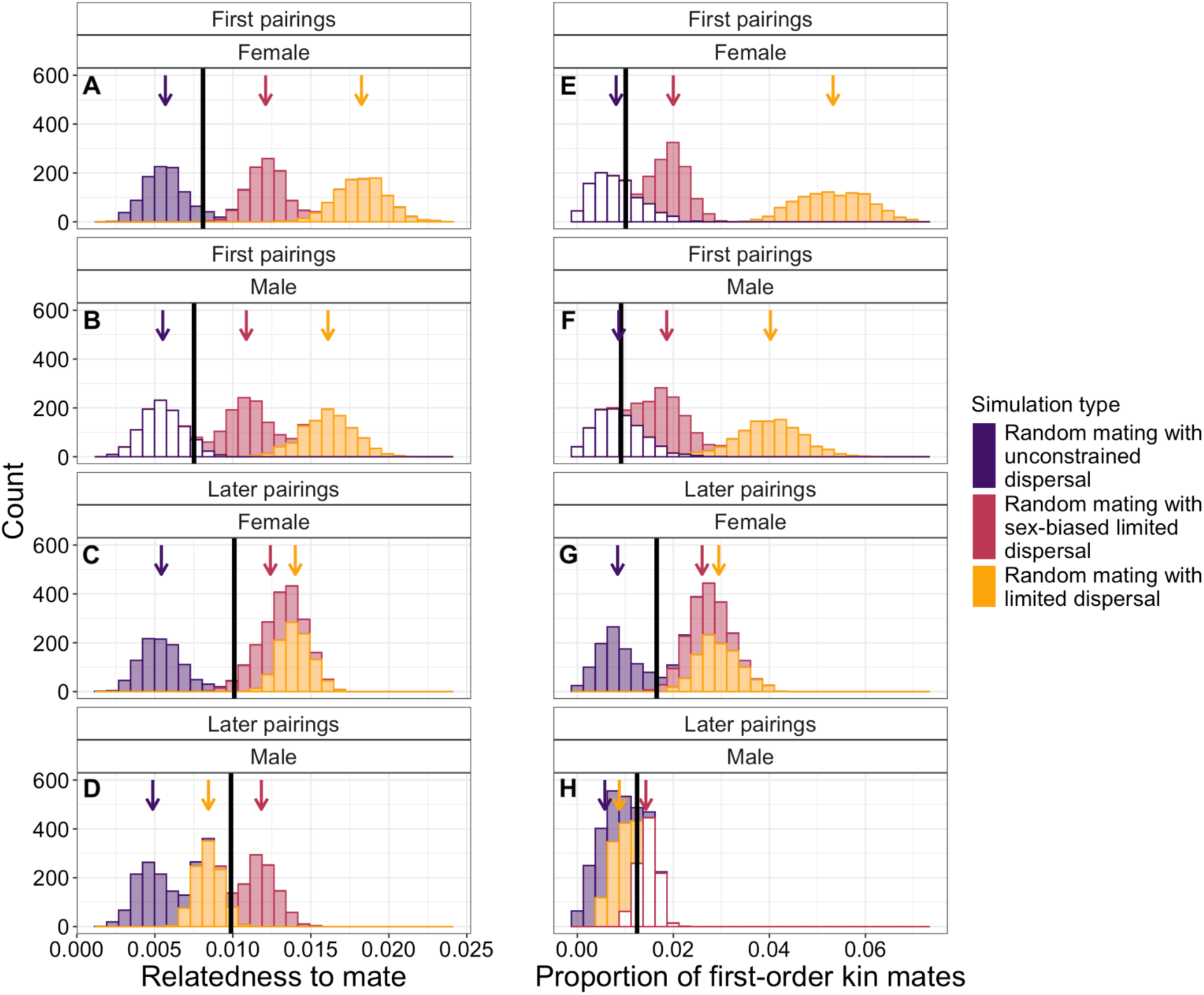
Relatedness to mate (A-D) and proportion of first-order kin mates (E-H) observed and expected under three different models of random mating in Florida scrub-jays at two life stages (first (A-B, E-F) and later pairings (C-D, G-H)) and for both sexes (male (B, F, D, H) and female (A, E, C, G)). The observed population means across all 32 years of our study period are indicated by black vertical lines. Histograms show the distributions of expected means under 1000 simulations of random mating in a scenario with unconstrained dispersal (purple), limited and sex-biased dispersal (red), and limited dispersal without sex-bias (yellow) (filled bars = distribution significantly different; unfilled bars = distribution not significantly different from the observed mean). Means of distributions are indicated by arrows.

### Fitness consequences of inbreeding versus inbreeding avoidance

*Breeder survival:* For both sexes, breeder survival was affected by distance moved for first pairing: farther distance moved was associated with higher mortality in the first four years after breeding onset (Males: *Z* = 2.16, *P* = 0.03; Females: *Z* = 3.36, *P* = 0.001), but the relationship was reversed in later years (Males: *Z* = -2.07, *P* = 0.04; Females: *Z* = -2.02, *P* = 0.04) (Fig. S9). Though the median breeding lifespan for both sexes was 4 years, 37% of individuals lived past their fourth year as a breeder (Males: 73 out of 197, Females: 45 out of 120 individuals). However, dispersal away from an established breeding territory in the event of later pairing did not impact breeder survival for either sex (Males: *Z* = -0.37, *P* = 0.71; Females: *Z* = 0.76, *P* = 0.45).

#### Lifetime reproductive success

Greater weighted kinship coefficients to mates over individuals’ lifetimes were associated with significantly fewer offspring that survived to adulthood (*i.e.*, LRS) for males (Fig. 2B, D; Tables 1-2), but not females (Tables 1-2; Fig. 2B, D). The few males who bred with closely related females for the majority of their lifetime had LRS close to zero (Fig. 2B), but we did not observe the same pattern in females. However, this biologically peculiar result may simply reflect tiny sample sizes, since only 6% of individuals in our dataset (11 males, 8 females out of 317 breeders) had weighted kinship coefficients to mates > 0.05. Females, but not males, who moved farther for later pairings had lower associated LRS, though the effect was only marginally significant (Table 2 Fig. 2C). After controlling for the significant negative effect of breeding lifespan—which is lower for individuals with longer natal dispersal distances—on LRS (Table 1), we did not find an additional effect of distance moved for first pairing on LRS for either sex (Table 1, Fig. 2A).

**Figure 2.**
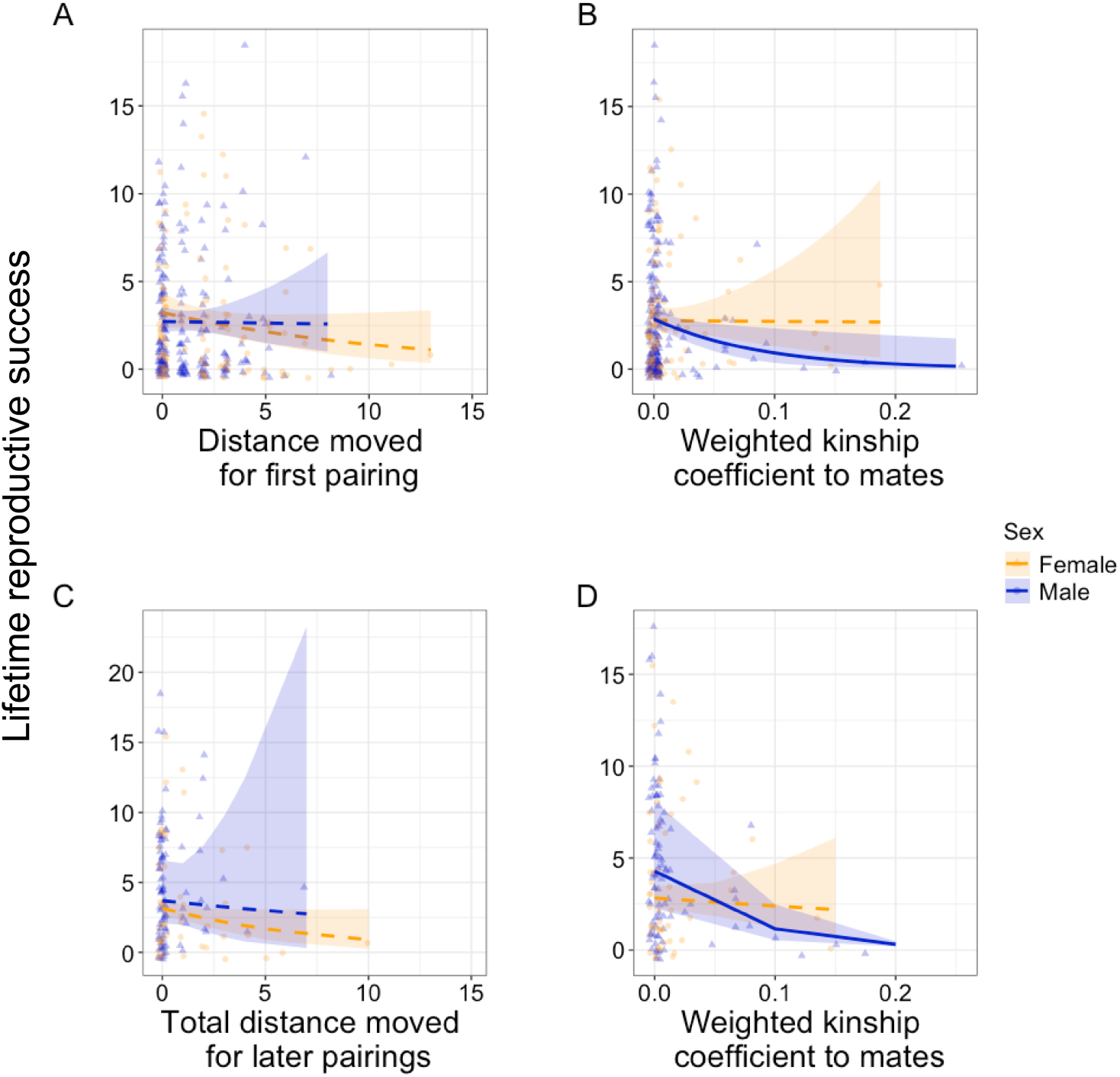
Predicted model fits investigating the effects (solid line = significant, dashed line = not significant; shaded area = 95% confidence interval) of (A) Distance moved for first pairing, and (B) Weighted kinship coefficients to mates on lifetime reproductive success (LRS) for Florida scrub-jay male (blue, triangles) and female (orange, circles) breeders with lifetime data between 1990 and 2021 (*N*: Males = 197, Females = 120). For the subset of breeders that switched mates at least once in their lifetime (*N*: Males = 91, Females = 50), predicted model fits investigating the effects of (C) Total distance moved for later pairings, and (D) Weighted kinship coefficients to mates on LRS. Female LRS showed a marginally significant negative association with greater total distance moved for later pairings (*P =* 0.06), while male LRS showed a significant negative association (*P* ≤ 0.05) with higher kinship to mates over their lifetime. Points are jittered both horizontally and vertically for better visual discrimination.

**Table 1.**
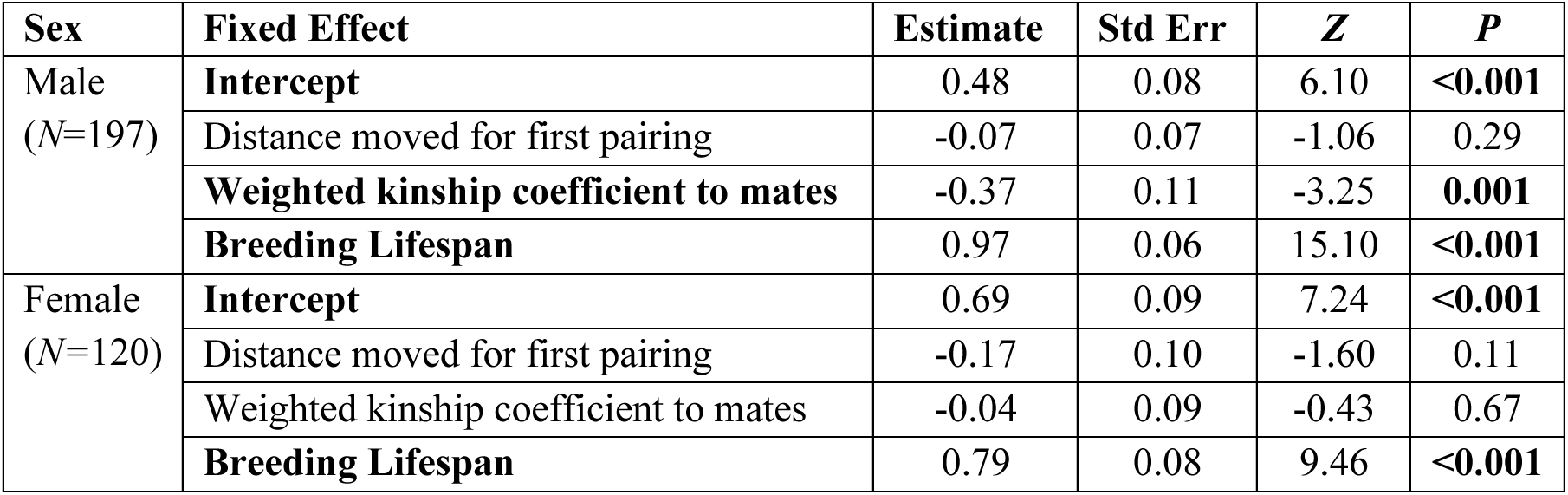
Results of generalized linear mixed models estimating the effect of distance moved for first pairings, relatedness to mate, and breeding lifespan on lifetime reproductive success (*i.e.*, number of offspring surviving to adulthood) of Florida scrub-jay breeders. Models were fit separately by sex using a negative binomial error distribution. Significant effects are in bold (P ≤ 0.05).

| Sex | Fixed Effect | Estimate | Std Err | Z | P |
| --- | --- | --- | --- | --- | --- |
| Male<br>(N=197) | **Intercept** | 0.48 | 0.08 | 6.10 | **<0.001** |
|  | Distance moved for first pairing | -0.07 | 0.07 | -1.06 | 0.29 |
|  | **Weighted kinship coefficient to mates** | -0.37 | 0.11 | -3.25 | **0.001** |
|  | **Breeding Lifespan** | 0.97 | 0.06 | 15.10 | **<0.001** |
| Female<br>(N=120) | **Intercept** | 0.69 | 0.09 | 7.24 | **<0.001** |
|  | Distance moved for first pairing | -0.17 | 0.10 | -1.60 | 0.11 |
|  | Weighted kinship coefficient to mates | -0.04 | 0.09 | -0.43 | 0.67 |
|  | **Breeding Lifespan** | 0.79 | 0.08 | 9.46 | **<0.001** |

**Table 2.** Results of generalized linear mixed models estimating the effect of total distance moved for later pairings, relatedness to mate, and breeding lifespan on lifetime reproductive success (*i.e.*, number of offspring surviving to adulthood) of Florida scrub-jay breeders who switched mates at least once in their lifetime. Models were fit separately by sex. For males we used a negative binomial error distribution, but for females we used a zero-inflated Poisson error distribution due to the small sample size and model convergence issues. Significant effects are in bold (P ≤ 0.05).

| Sex | Model component | Fixed effect | Estimate | Std Err | Z | P |
| --- | --- | --- | --- | --- | --- | --- |
| Male<br>(N=91) | - | **Intercept** | 1.29 | 0.08 | 17.00 | **<0.001** |
|  |  | Total distance moved for later pairings | -0.04 | 0.06 | -0.72 | 0.47 |
|  |  | **Weighted kinship coefficient to mates** | -0.42 | 0.11 | -3.72 | **<0.001** |
|  |  | **Breeding Lifespan** | 0.58 | 0.07 | 8.76 | **<0.001** |
| Female<br>(N=50) | Count model | **Intercept** | 1.01 | 0.10 | 10.15 | **<0.001** |
|  |  | Total distance moved for later pairings | -0.24 | 0.13 | -1.88 | 0.06 |
|  |  | Weighted kinship coefficient to mates | -0.05 | 0.11 | -0.45 | 0.65 |
|  |  | **Breeding Lifespan** | 0.83 | 0.08 | 10.49 | **<0.001** |
|  | Zero-inflation model | Intercept | -6.20 | 3.79 | -1.64 | 0.10 |
|  |  | Breeding Lifespan | 2.12 | 2.00 | 1.06 | 0.29 |

### Degree of inbreeding avoidance via mate choice

We found that observed mean relatedness to mates was higher than expected by chance under a model of unconstrained dispersal with random mating (scenario i) for both sexes at all life stages, except males for first pairings where the difference approached significance (Males, First pairings: *P* = 0.07; Females, First pairings: *P* = 0.05; Males, Later pairings: *P* = 0.001; Females, Later pairings: *P* = 0.003; Fig. 1A-D). However, observed mean relatedness to mates remained significantly lower across all sexes and life stages than expected under the limited, sex-biased dispersal with random mating model (scenario ii), indicating the existence of inbreeding avoidance via mate choice (Males, First pairings: *P* = 0.005; Females, First pairings: *P* < 0.001; Males, Later pairings: *P* = 0.03; Females, Later pairings: *P* = 0.04; Fig. 1A-D).

We also found that observed proportions of pairings between first-order kin were not significantly different from those expected under the unconstrained dispersal with random mating model (scenario i) (Males: *P* = 0.43, Females: *P* = 0.32, Fig. 1E-F) but were significantly lower than those expected under the random mating with limited, sex-biased dispersal model (scenario ii) in both sexes for first pairings (Males: *P* = 0.01, Females: *P* < 0.001, Fig. 1E-F). Conversely, for later pairings, the observed proportions of first-order kin were significantly higher than expected under the unconstrained dispersal with random mating model (scenario i) (Males: *P* = 0.05; Females: *P* = 0.04, Fig. 1G-H), and, for males, not significantly different from expectations under the random mating with limited, sex-biased dispersal model (scenario ii) (Males: *P* = 0.28, Females: *P* = 0.008, Fig. 1G-H). For second- and third-order kin mates, patterns were more variable across sexes and dispersal types (Fig. S10). Overall, both sexes exhibited higher proportions of non-kin mates than expected under the random mating with limited, sex-biased dispersal model (scenario ii) for first pairings, but not for later pairings (Males, First pairings: *P* = 0.009, Females, First pairings: *P* = 0.04, Males, Later pairings: *P* = 0.10, Females, Later pairings: *P* = 0.17; Fig. S10).

Relatedness to their mate had no effect on likelihood of mate switching for either sex at either life stage (Males, First pairing: *Z* = 1.41, *P* = 0.16; Males, Later pairing: *Z* = 0.42, *P* = 0.68; Females, First pairing: *Z* = -0.86, *P* = 0.39; Females, Later pairing: *Z* = 0.76, *P* = 0.45), nor was mate switching more likely in cases of pairing with first-order relatives, though the effect was marginally significant for males during first pairings (Males, First pairing: *Z* = 1.87, *P* = 0.06; Males, Later pairing: *Z* = 0.22, *P* = 0.83; Females, First pairing: No instances of mate switching when paired with a first-order relative; Females, Later pairing: *Z* = 0.54, *P* = 0.59).

## Discussion

We empirically tested the theoretical prediction that inbreeding avoidance is not adaptive under all conditions (Kokko and Ots 2006; Waser et al. 1986). While studies in other taxa, such as plants and fungi (Karron et al. 2012; Nieuwenhuis and James 2016), have shown that inbreeding tolerance, or even preference, can be adaptive when the fitness costs of inbreeding avoidance outweigh the fitness costs of inbreeding, inbreeding in animals is still considered universally maladaptive (de Boer et al. 2021). However, investigating the life history traits and relative fitness outcomes underlying the evolution of inbreeding avoidance in animals is challenging since it requires detailed observational data across individuals’ full lifetimes. Here, we leverage a unique dataset from a long-lived avian species with a low propensity for and distance of dispersal (*i.e.*, limited dispersal) (Suh et al. 2020) experiencing inbreeding depression (Chen et al. 2016). First, we showed that limited dispersal creates the scope for the evolution of inbreeding avoidance by significantly increasing inbreeding risk. Though this risk is mitigated by sex-biased dispersal, the effect varies by life stage and does not eliminate inbreeding risk. Second, we found that both dispersing farther and pairing with close relatives are associated with lower lifetime fitness, suggesting that the cost of inbreeding avoidance via dispersal is equivalent to or higher than the cost of inbreeding, except when breeding with first-order kin. Third, we found that the inbreeding risk introduced by dispersal constraints is tolerated to a significant extent, and once paired with relatives, individuals are not more likely to switch mates to avoid inbreeding. However, within the dispersal constraints, we found that individuals actively avoid breeding with first-order kin. Taken together, our results suggest that categorical inbreeding avoidance need not be the norm. Even in a population with inbreeding depression, some degree of inbreeding tolerance can exist, and a more nuanced investigation of the conditions that may select for inbreeding tolerance versus avoidance is recommended in future studies (Kokko and Ots 2006).

### Limited dispersal creates the scope for the evolution of inbreeding avoidance by increasing inbreeding risk

We found that a key life-history trait—limited dispersal—contributes significantly to elevated inbreeding risk, creating a condition potentially favoring the evolution of inbreeding avoidance (Duthie and Reid 2016). Although elevated risk of inbreeding and associated inbreeding depression might be expected to select for longer dispersal distances, such dispersal limitations and associated inbreeding risk are surprisingly common in both social and asocial species across taxa (Bell et al. 2024; Ceresa et al. 2024; Comte and Olden 2018; Costantini et al. 2007; De Ro et al. 2021; Keller et al. 1994; Lloyd et al. 2018; Lubin et al. 2009). Limited dispersal can lead to a variety of fitness benefits: reproductive assurance (Moeller 2006), access to resources and protection from predators due to prior local experience (Pärt 1994; Suh et al. 2020), lower competition (Green and Hatchwell 2018; Kokko and Ekman 2002), and lower physiological demands (Ashman and Schoen 1996; Maag et al. 2019). We also separately considered instances of all subsequent pairings with new mates. We found that high site fidelity, *i.e.,* limited movement away from the breeding territory for later pairings, also significantly increased inbreeding risk. While studies have rarely examined the contribution of dispersal patterns in later life stages to inbreeding risk, spatiotemporal consistency in rutting behavior in red deer (*Cervus elaphus*) has been shown to result in inbreeding by increasing the chances of males mating with females of the same matriline (Stopher et al. 2012). Thus, neglecting later stages of life excludes demographic processes that can significantly contribute to inbreeding, especially in long-lived species.

Theoretical work suggests that in sexually reproducing species, sufficiently higher propensity and distance of dispersal in only one sex can effectively mitigate inbreeding risk (Li and Kokko 2019; Perrin and Mazalov 1999). However, to our knowledge the extent to which sex-biased dispersal mitigates inbreeding risk has only been quantified in one other system (Lebigre et al. 2010). We isolated the effect of sex-biased dispersal on mitigating inbreeding risk *in silico*. We found that expected relatedness to mates was significantly lower under a random mating scenario with limited, but sex-biased, dispersal compared to a scenario with limited dispersal without sex-bias during the first pairing stage. However, our data show that Florida scrub-jay females still show considerable philopatry at the first pairing stage, only dispersing on average ∼2 territories away for first pairings. The likelihood of encountering a closely related potential mate is thus still significantly higher within the average dispersal distance of both sexes than at farther dispersal distances. These results add to a growing body of literature in social vertebrates where inbreeding risk persists despite sex-biased dispersal due to the favorable fitness benefit of remaining in familiar surroundings, in the same social group as kin, or close to kin (Daniels and Walters 2000b; Kingma et al. 2017; Morrison et al. 2023; Riehl and Stern 2015). Even in asocial species, other life history traits, such as overlapping generations and iteroparity (Blyton et al. 2015), or anthropogenic impacts such as habitat fragmentation (Athrey et al. 2012) can increase inbreeding risk despite sex-biased dispersal. Thus, the degree to which sex-biased dispersal leads to inbreeding avoidance is highly context dependent.

At later pairing stages, we found that males were twice as likely as females to remain on their breeding territory. Such sex bias in site fidelity is widespread in territorial species, but which sex is more dispersive can depend on mating systems, reproductive investment and incest avoidance (Berteaux and Boutin 2000; Clarke et al. 1997; Packer 1979; Packer and Pusey 1987). Contrary to our predictions, however, removing the sex bias in distance moved for later pairings—*i.e.*, relaxing the strong skew in males towards site fidelity—decreased, rather than increased, inbreeding risk in males by 50% (though this decrease is countered by a 17% increase in inbreeding risk for females). This result suggests that the role of sex-biased dispersal in mitigating inbreeding risk may be highly context dependent. While studies of inbreeding avoidance largely focus on sex bias in natal dispersal (*e.g.*, Morrison et al. 2023; Schiegg et al. 2006), variation in dispersal propensity and distance between sexes for later pairings should be considered for a more thorough understanding of the processes that create inbreeding risk (Pusey 1987).

### Fitness trade-offs can favor inbreeding tolerance

We found that lifetime fitness costs of inbreeding do not outweigh the fitness benefits of limited dispersal, suggesting that strict inbreeding avoidance is not necessarily adaptive in this species. These relative fitness outcomes may instead result in a degree of inbreeding tolerance (Kokko and Ots 2006; Waser et al. 1986). Florida scrub-jays experienced fitness costs with longer distances moved at all life stages: natal dispersal distance is negatively associated with breeder survival (*i.e.*, breeding lifespan)—which we found has the strongest effect on lifetime reproductive success (LRS), a relationship also shown in previous studies (Fitzpatrick et al. 1999; Summers et al. 2024)—and distance moved for later pairings showed a trend towards negative association with LRS, though only in females. Note that the small sample size for females with later pairings constrained our use of the best-suited model. However, our analysis likely underestimates the cost of moving farther distances, since it does not include individuals who die in transit or otherwise fail to establish as breeders as well as dispersers out of the study tract (Bonte et al. 2012). We note that these effects may be due to lower-quality individuals dispersing farther and losing competition for mates or breeding territories rather than a cost of dispersal itself (Fuirst et al. 2023; Hui et al. 2012). Lifetime fitness costs of inbreeding were evident only in males, but as mentioned above, this result must be interpreted with caution since individuals that breed with closely related mates, especially for a high proportion of their lifetimes, are extremely rare in our population. Though survival rates are equal between the sexes, females may be able to compensate for a period of low reproductive success in their lifetime via strategies not available to males (Woolfenden and Fitzpatrick 1987), such as varying reproductive investment (Harris and Uller 2009) or increased choosiness (Gabor and Halliday 1997).

LRS only approached zero for the few males that bred with closely related females for most of their lifetimes, which matches the pattern of reduced hatching success in closely related pairs shown previously in this species (Chen et al. 2016). Thus, inbreeding costs may be elevated when breeding with kin, but breeding with more distant relatives appears not to be costly enough to outweigh the benefits of limited dispersal (Kokko and Ots 2006). This pattern could be a result of purging, where initial inbreeding leads to the rapid loss of strongly deleterious and lethal variants from the population such that the fitness costs of inbreeding lessen over generations (Hedrick 1994). Though studies of reproductive success of breeders engaging in modest levels of inbreeding are few, they show a similar pattern of little to no effect on annual or lifetime reproductive success (Frappier-Lecomte et al. 2025; Rowley et al. 1986; Van de Casteele et al. 2003; Willoughby et al. 2019). Fitness effects instead manifest over the inbred offspring’s lifetime (Chen et al. 2016; Husband and Schemske 1996), and the degree of inbreeding depression is highly variable within and across species (Angeloni et al. 2011; Vega-Trejo et al. 2022). In sum, our results show how the fitness benefits of a life history trait that increases inbreeding risk can outweigh the costs of inbreeding, contributing to inbreeding tolerance instead of avoidance. Further, a low degree of inbreeding could also result in inclusive fitness (Kokko and Ots 2006) and guard against outbreeding depression (Bateson 1983). Empirical evidence for such benefits of inbreeding is limited (Bateson 1982), and future studies should investigate these possibilities in both this and other study systems.

### Inbreeding avoidance depends on relatedness and life stage

In line with theoretical predictions (Bengtsson 1978; Kokko and Ots 2006) we found that while limited dispersal creates a condition favoring the evolution of inbreeding avoidance, Florida scrub-jays tolerate the modest level of inbreeding risk introduced by limited dispersal. This pattern may be explained by the relatively lower fitness costs of inbreeding compared to those of inbreeding avoidance by dispersal. Additionally, in line with our finding that LRS approaches zero for males when their mates are consistently first-order kin, we found evidence of kin avoidance, specifically of first-order kin, within the observed dispersal constraints for first pairings. As a recent meta-analysis suggests, such avoidance of close kin mates may explain the prevailing view that inbreeding avoidance is always adaptive (de Boer et al. 2021), since studies largely test for choice between first- or second-order kin and non-kin only (Hidalgo Aranzamendi et al. 2016; Krause et al. 2012; Leedale et al. 2020; Morrison et al. 2023; Välimäki et al. 2011). Exploring inbreeding tolerance across a continuum of relatedness to mates, and within the context of life history constraints—such as limited dispersal—may reveal a greater prevalence of inbreeding tolerance in animal systems (Kokko and Ots 2006). In plants, studies have shown that a mix of dispersal syndromes can evolve when dispersal is not always the optimal strategy, similarly resulting in a degree of inbreeding tolerance (Cheptou and Massol 2009; Hidalgo et al. 2016).

Mechanisms underlying kin avoidance could include phenotype-matching using auditory, visual, or olfactory cues (Bateson 1982; Bonadonna and Sanz-Aguilar 2012; Krause et al. 2012; Sharp et al. 2005) or prior experience and social familiarity. Since Florida scrub-jays live in cooperative breeding groups where the majority of helpers are offspring from previous broods that delay dispersal for one or more years to act as helpers (Woolfenden and Fitzpatrick 1984), first-order kin grow up having ample opportunity to build familiarity with close kin, which can facilitate kin recognition (Komdeur and Hatchwell 1999). In fact, kin-directed helping may additionally select for the evolution of kin recognition via either mechanism (Nichols 2017). Similar patterns of close-kin avoidance have been shown in a range of social species in taxa ranging from insects (Lizé et al. 2014) to fish (Hain and Neff 2007), birds (Cockburn et al. 2003; Hatchwell et al. 2000; Haydock et al. 2001), and mammals (Galezo et al. 2022; Morrison et al. 2023).

Interestingly, we found that the pattern of kin tolerance was stronger, with the effect size an order of magnitude higher, for later pairings than first pairings. Additionally, for later pairings, both sexes paired with first-order kin more than expected by chance and as expected under a random mating scenario within their observed dispersal distances. This result further suggests that, with a shift in fitness tradeoffs, inbreeding tolerance may increase for Florida scrub-jays at later life stages. Losing a mate can substantially weaken the remaining mate’s ability to retain its territory, and thus its breeding status as well as familiar and cached food resources (Woolfenden and Fitzpatrick 1984, 1987). Thus, the fitness benefits of site fidelity may outweigh the fitness benefits of inbreeding avoidance for established breeders. Such benefits of site fidelity (Beck et al. 2020; Piper et al. 2023; Schwartz et al. 2007), and associated increase in site fidelity with age (Cameron et al. 2007; Piper et al. 2018; Switzer 1997), have been documented in a variety of territorial species. Additionally, a variety of species also show changes in mate choice behavior with age (Edward and Chapman 2013; Gabor and Halliday 1997; Pitcher et al. 2003). Our results suggest that the association between site fidelity and mate choice merits further study. More broadly, future studies should investigate how the relative costs and benefits of inbreeding tolerance shift with age, experience, and successive mate choice in the wide range of taxa that experience more than one mate choice opportunity in their lifetimes (Edward and Chapman 2013; Foerster et al. 2006; Gabor and Halliday 1997; Marshall and Folsom 1991; Pitcher et al. 2003).

Finally, though mate switching is a common inbreeding avoidance strategy across animal taxa (Hatchwell et al. 2000; Cockburn et al. 2003; Hidalgo Aranzamendi et al. 2016), we did not find a consistent pattern. Only males in first pairings were marginally more likely to switch mates when paired with first-order kin females. Pairs consisting of close relatives are rare in our study population, with only 4% of all breeding pairs resulting in consanguineous matings and only 1.2% resulting in matings between first-order relatives. Kin avoidance may reduce the likelihood of first-order kin pairs forming in the first place. Further, mate switching may prove costly in terms of territory and breeding status loss, especially for females, in our study population where breeding vacancies are infrequent (Woolfenden and Fitzpatrick 1984), a substantial fitness cost that may not exist in populations of other pair-bonding species where mate switching is a more common inbreeding avoidance strategy (Cockburn et al. 2003; Hatchwell et al. 2000; Hidalgo Aranzamendi et al. 2016; Otter and Ratcliffe 1996).

## Conclusions

We studied how dispersal and mate choice decisions contribute to inbreeding risk and the conditions underlying the evolution of inbreeding avoidance in a natural population experiencing inbreeding depression. We found that a key life-history trait—limited dispersal—increased inbreeding risk and created the scope for the evolution of inbreeding avoidance, but the lifetime fitness costs of inbreeding did not exceed those of dispersing farther, except for individuals that bred with first-order kin. Accordingly, we found that individuals only avoided breeding with first-order kin within dispersal constraints, with the degree of inbreeding avoidance varying with both sex and life stage. Taken together, these results corroborate theoretical predictions that the evolution of inbreeding avoidance is context dependent and not always adaptive (Bengtsson 1978; Duthie and Reid 2016; Kokko and Ots 2006; Waser et al. 1986). Future observational and experimental studies across species should question the assumption that inbreeding avoidance is always adaptive and consider the specific fitness outcomes across traits such as sex and life stage to better understand when inbreeding avoidance evolves, and when it does not.

## Acknowledgments

We thank Archbold Biological Station and its staff for continued support and management of the study area; all interns, students, and postdocs who have worked on collecting and managing the long-term data; the Cornell Statistical Consulting Unit for statistical support; V. Sclater for GIS support; and B. Kirz, L. Grieves, F. Romero, C. Rushworth, D. Seidman, S. Siller, J. Summers, D. Tian, and M. Webster for invaluable feedback on previous versions of this manuscript.

## Funding

This work was supported by National Science Foundation (NSF) Grants DEB0855879 and DEB1257628 and by the Cornell Lab of Ornithology Athena Fund. S.S.S was supported by NIH grant 1R35GM133412 to N.C. and NSF Postdoctoral Research Fellowship in Biology 2305705.

## Notes

### Competing Interest Statement

The authors have declared no competing interest.

### Summary of Updates

We revised the writing for broader appeal

