## Supplementary Material for "Lifetime fitness benefits of short-distance dispersal are associated with inbreeding tolerance despite multiple inbreeding avoidance mechanisms"

This PDF file includes:
Supplementary Text
Figures S1-S14
Tables S1-S5
References

**Supplementary Text**

*Descriptive statistics for late pairings*

In 197 out of 848 (23%) instances of later pairings, the focal individual (62% of whom were females) moved one or more territories away from its established breeding territory. Only about half of all mates in later pairings were naïve breeders for both sexes (Males: 218 out of 427 [51%], Females: 213 out of 421 [51%] instances). Thus, not all later pairings were a result of subordinates moving in to fill a breeding vacancy—nearly half of such cases were a result of the mate also re-pairing because of death of their previous mate or mate switching. However, for breeders that retained their breeding territory for later pairings, ≥ 90% of new mates were from ≤ 2 territories away (Fig. S6). Furthermore, we found that out of all later pairing events observed, 18% were due to mate switching, not mate death (Males = 33 out of 187; Females = 31 out of 169 pairings).

*Territory of origin for calculating distance moved for pairing*

The previous territory—*i.e.*, territory of origin—for any movement associated with a new pairing was identified as the territory where the individual was seen or caught the previous year between March and June (breeding season), and, if the bird was not seen within that time duration, between July to December (post-breeding season) since dispersal peaks at the start and end of the breeding season (Suh et al. 2020). For "staging" individuals that eventually became breeders—*i.e.*, individuals that initially dispersed to become helpers at a non-natal territory—we used the distance from their staging territory to their breeding territory to as distance moved for first pairing. However, using the natal territory as territory of origin did not change our results (Fig. S11, Table S4). Additionally, when two experienced breeders formed a new breeding pair, their breeding territories in the previous breeding season were identified as their previous territories. We also accounted for territory vacancies filled throughout the year. In instances when a first-time breeder formed a new pair with an experienced breeder who retained its breeding territory, we noted the month preceding the last month that the previous mate was seen on the territory. We then identified the territory of origin for the first-time breeder as the territory in which it was seen in that month.

*Manual, case-by-case quantification of number of territories crossed*

Florida-Scrub Jay territories at Archbold Biological Station are assigned a unique name based on their location. While territory boundaries shift stochastically from year to year, territories retain the same name if the general location and at least one of the two dominant breeders remain the same. Thus, for cases where our automated calculations indicated that an individual moved at least one or more territories away, but the territory name associated with them remained the same, we (J.W.F., S.B., and S.S.S.) manually assigned distance moved on a case-by-case basis ($N$ = 88 out of 1669 [5%]). For territories that were roughly in the same location as the previous year, with the territories around it also retaining roughly the same orientations and arrangements, we assigned a distance moved of 0 territories away (Fig. S12A). However, if the territory location had changed such that its orientation with respect to surrounding territories had changed, then we assigned a distance moved of ≥ 1 territory away depending on how many territories were between its previous and its new location (Fig. S12B).

*Sex-bias in distance moved at different life stages*

To examine sex bias in distance moved for first and later pairings, we ran generalized linear mixed models (GLMMs) with distance moved as the dependent variable, sex as the fixed effect, and a random effect of year. A random effect of individual ID was also included in the model for distance moved for later pairings to account for individuals who re-paired multiple times in their lifetime. The models had a negative binomial error distribution and were run separately by pairing type using the function *glmer.nb* in the R package *MASS* (Venables and Ripley 2002). We found that though females moved farther than males for pairing at both life stages, the sex-bias in distance moved was higher for first pairings (Males = 1.05 ± 1.53, Females = 1.90 ± 2.50 territories crossed; $Z = 5.86$, $P < 0.001$) compared to later pairings (Males = 0.30 ± 0.78, Females = 0.56 ± 1.17 territories crossed; $Z = 3.64$, $P < 0.001$) (Fig. S1). Overall, compared to an a random mating scenario with unconstrained dispersal, both sexes moved significantly shorter distances at both life stages within the study tract ($P \leq 0.05$ for all, Fig. S13).

*Relatedness to potential mates based on geographical distance*

To examine how relatedness to potential mates varies with distance, for each year we paired individuals of the focal sex with all available individuals of the opposite sex in a given year and calculated the potential dispersal distance and kinship coefficient to mate as described in the *Main Text* (*N*: Males, First pairing = 426; Males, Later pairing = 241; Females, First pairing = 395; Females, Later pairing = 251 individuals). We classified potential mates to whom the kinship coefficient of the focal individual was ≥ 0.0625 as close relatives (at least as related as first cousins, Fig. S7). We then fit generalized linear mixed models (GLMMs) with whether the potential mate was a close relative (1 = yes, 0 = no) as the dependent variable, a binomial error structure, distance to mate as a fixed effect, and year and focal individual ID as random effects. We fit separate models for each sex and pairing type (*N*: Males, First pairing = 12,111; Males, Later pairing = 11,121; Females, First pairing = 10,800; Females, Later pairing = 10,464 possible pairings). We found that both sexes were more closely related to potential opposite-sex mates at shorter geographical distances on both their natal and breeding territories (Males, Natal: $Z = -11.37$, $P < 0.001$; Females, Natal: $Z = -14.53$, $P < 0.001$; Males, Breeding: $Z = -8.10$, $P < 0.001$; Females, Breeding: $Z = -10.07$, $P < 0.001$; Fig. S8).

*Using distance moved from natal to breeding territory as distance moved for first pairing for all breeders, including those that "staged"*

We identified individuals as "stagers" if they were observed on a territory other than their natal territory in the year before becoming a breeder for the first time. Consistent with a previous estimate that 28% of Florida scrub-jays stage post-fledging (Suh et al. 2022), we found that 26% (169 out of 659) of the breeders with known natal territories in our dataset staged. Data from Suh et al. 2022 show that the median number of staging years is two for both sexes (Table S5). We found that distance moved from staging to breeding territory (mean ± SD = 0.91 ± 1.41 territories crossed) is significantly lower than distance moved from natal to breeding territory (mean ± SD = 3.45 ± 3.30 territories crossed) (Mann-Whitney Test: $N = 148$, $U = 275.5$, $P < 0.001$). Here, we calculated distance moved from natal to first breeding territory for all breeders with known natal territories ($N = 659$) as the number of territories crossed in a straight line from the centroid of the natal to the centroid of the breeding territory (see *Main Text: Methods* for more detail). However, using distance from natal to breeding territory did not change the results of our models evaluating the effect of distance moved for first pairing, weighted kinship coefficient to mates, and breeding lifetime on lifetime reproductive success for either sex (Table S4), likely since only 66 out of 317 (21%) individuals with complete lifetime fitness had staged before becoming breeders, and distance travelled calculated as distance from territory occupied in the previous year to breeding territory and distance from natal territory to breeding territory were highly correlated (Pearson's $r = 0.70$, Fig. S11). Note that the correlation coefficient is not 1 even for individuals who did not stage because territory boundaries shift from year to year, so the number of territories crossed as calculated from the natal territory in the year the individual was born could be different than the number of territories crossed as calculated from that same territory in the year the individual dispersed to become a breeder.

*Determining time intervals for stratified Cox proportional hazards models*

We found that fitting Cox proportional hazard models with distance moved for first pairing as a fixed effect violated the proportional hazards assumption for both sexes (Males: $\chi^2 = 4.33$, df = 1, $P = 0.04$; Females: $\chi^2 = 4.21$, df = 1, $P = 0.04$), suggesting that the effect of distance moved for first pairing on the hazard rate varies over time. To account for this time-varying effect, we fit stratified Cox proportional hazards models with two time intervals: ≤ 4 years and > 4 years from the start of breeding (Zhang et al. 2018). We stratified the data at 4 years post breeding onset because the time-dependent coefficient showed a more negative trend in later years (Fig. S3). To verify this cut time point, we also fit proportional hazards models for each sex with a continuous, time-varying effect of distance moved for first pairing using the *timecox* function in the R package "timereg" (Scheike and Zhang 2011) and 1000 simulations. We found that the effect of distance moved for first pairing has an inflection point between 4 and 5 years post breeding onset for both sexes (Fig. S9). Thus, we stratified our data into ≤ 4 years and > 4 years post breeding onset and fit stratified Cox proportional hazards models. Stratifying data into time intervals with cut time points of 5-, 6-, and 7-years post breeding onset did not qualitatively change our results (Table S1). The upper quartiles of breeding lifespan were 6 years for males and 7 years for females (Fig. S14). The proportional hazard assumption for the model examining the effect of site fidelity on breeder survival was met without needing to incorporate a time-varying effect (Males: $\chi^2 = 1.17$, df = 1, $P = 0.28$; Females: $\chi^2 = 0.27$, df = 1, $P = 0.61$).

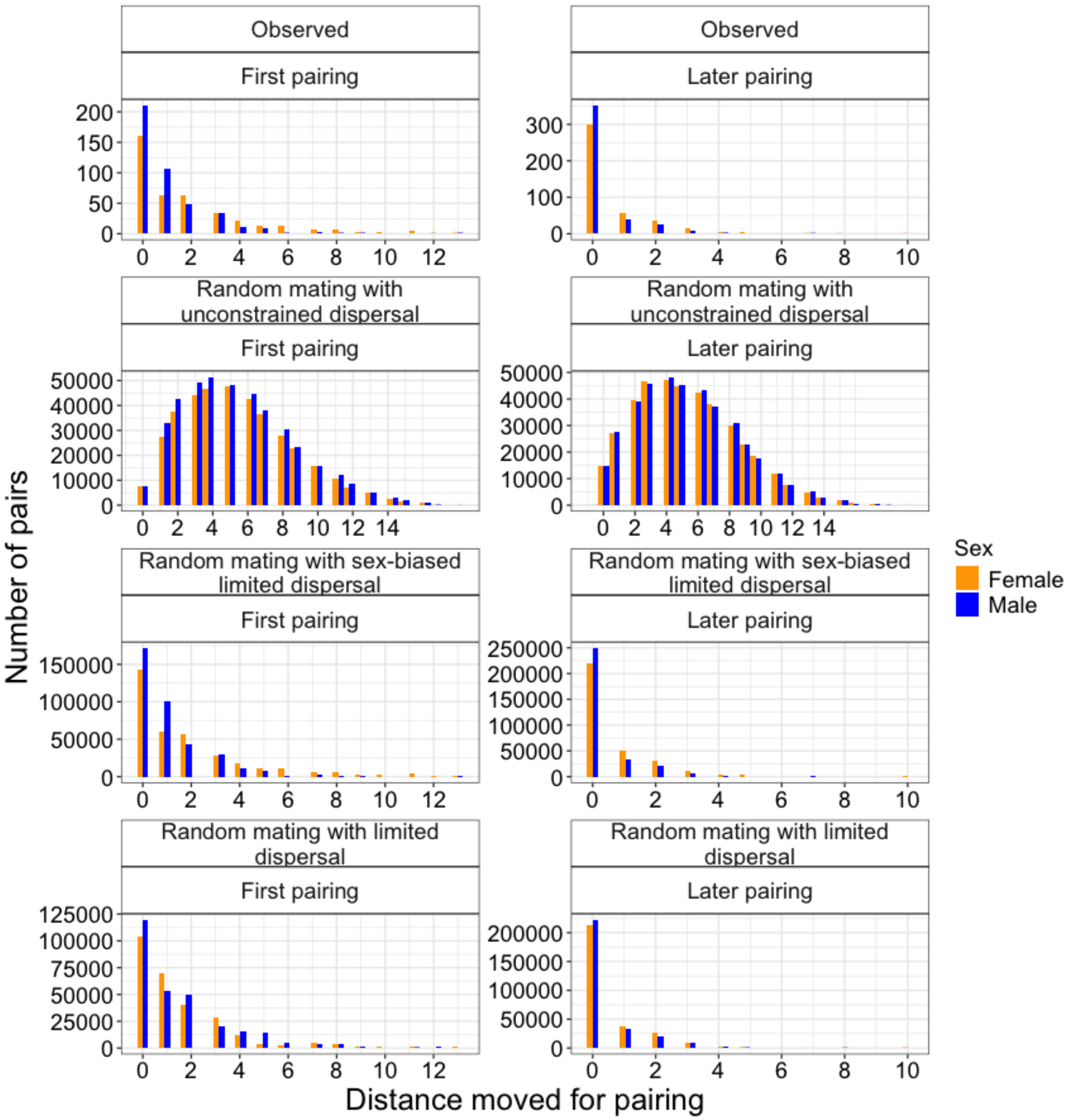


**Figure S1.** Histograms showing distributions of distances moved for first and later pairings observed in our study population and simulated in our random mating with (i) unrestricted dispersal, (ii) limited, sex-biased, and (iii) limited dispersal but no sex-bias models.

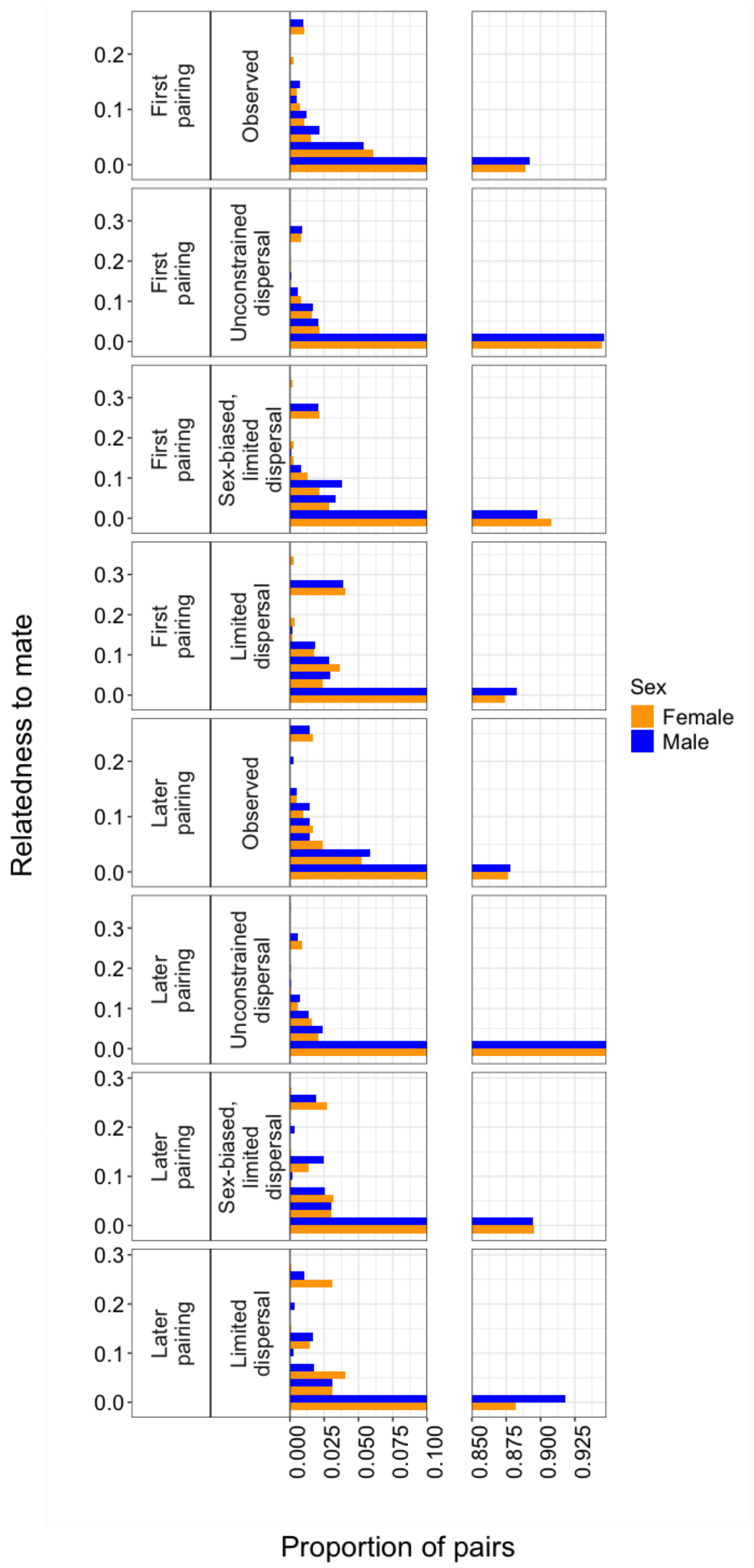


**Figure S2.** Histograms showing distributions of relatedness to mate observed in our study population and simulated in our random mating with (i) unrestricted dispersal, (ii) limited, sex-biased, and (iii) limited dispersal but no sex-bias models.

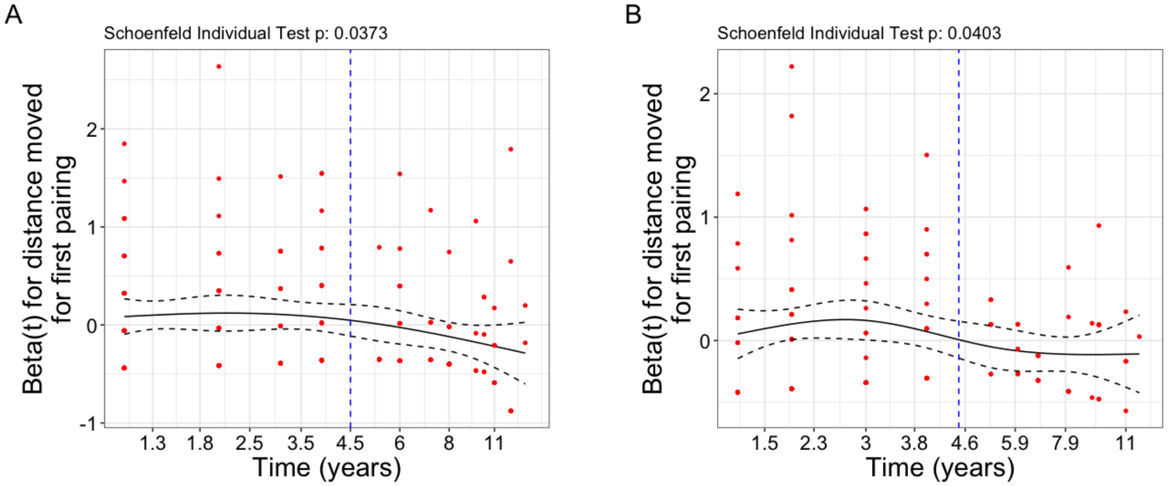


**Figure S3.** Plots showing how the time-dependent coefficient of distance moved for first pairing (y-axis) changes with time from breeding onset (x-axis) for (A) male, and (B) female Florida scrub-jays when fitting Cox proportional hazards models to estimate how distance moved impacts breeder survival. Between 4- and 5-years post breeding onset (*i.e.*, to the left and right of the blue dotted line), the effect of distance moved for first pairing on survival changes sign. Thus, the proportional hazard assumption is violated for both sexes (Males: $\chi^2 = 4.33$, df = 1, $P = 0.04$; Females: $\chi^2 = 4.21$, df = 1, $P = 0.04$).

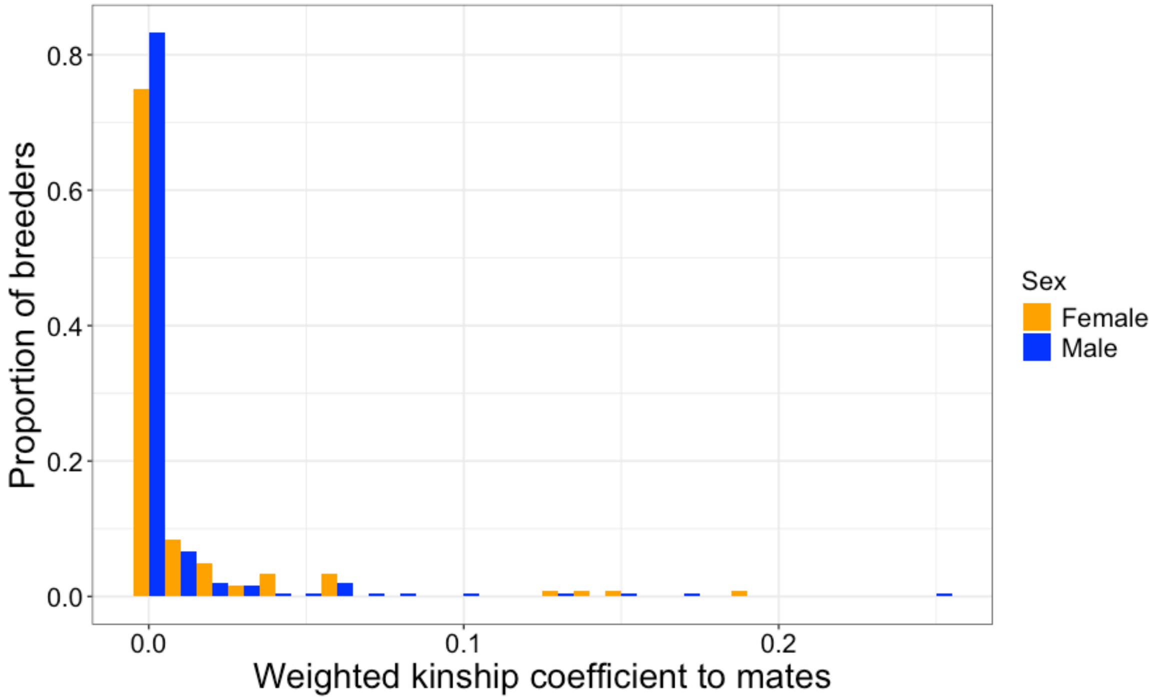


**Figure S4.** The distribution of weighted kinship coefficient to mates over the lifespan of female (orange) and male (blue) breeding Florida scrub-jays. Weighted kinship coefficient was calculated by multiplying the kinship coefficient of the focal individual to each of their mates by the number of years they were paired with that mate and summing the resulting values. Only 6% (19 out of 317) of individuals had a weighted kinship coefficient to mates > 0.05.

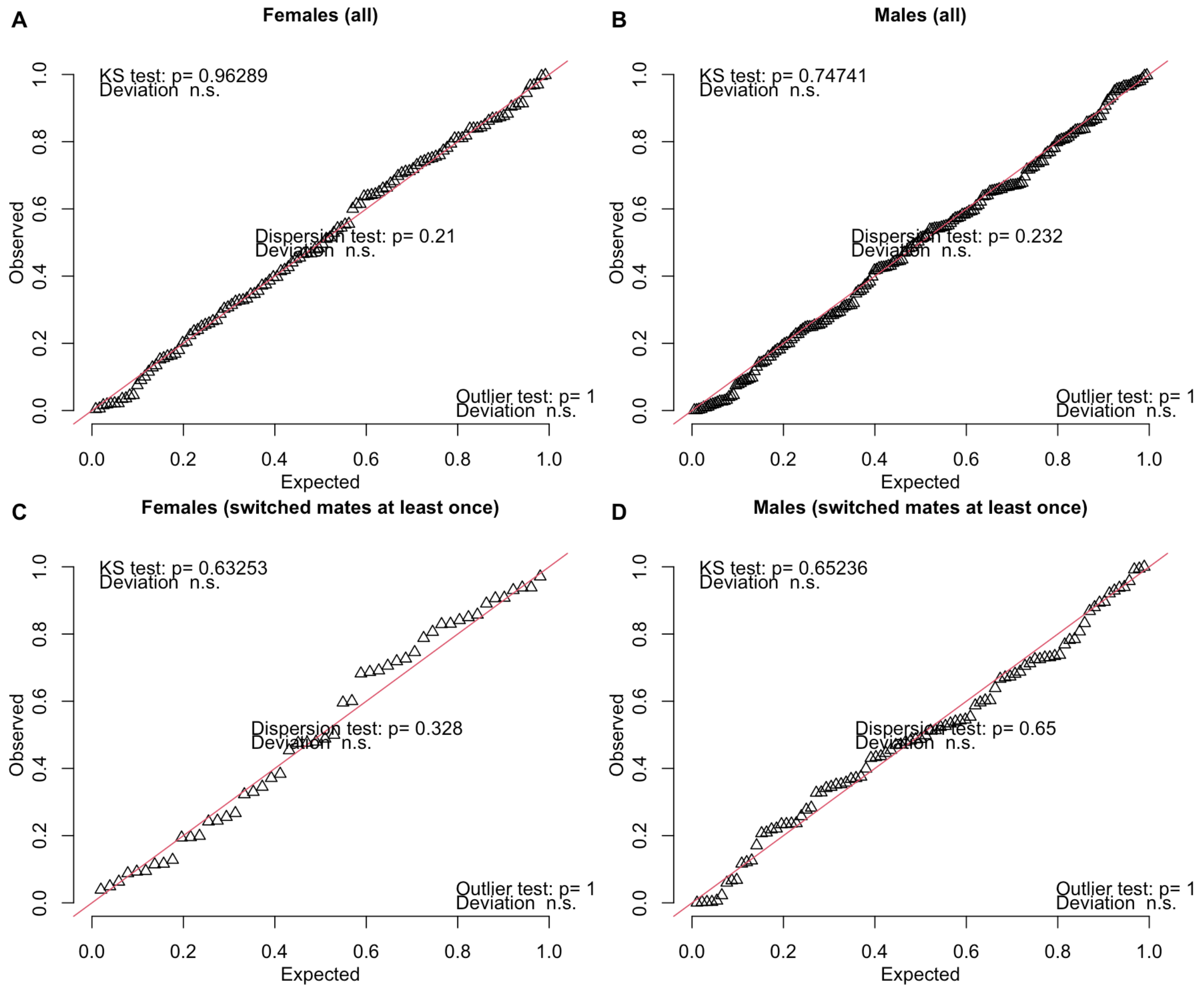


**Figure S5.** Quantile-quantile plots of residuals and additional tests of fit from models estimating effects on LRS of distance moved for first pairings, relatedness to mate, and breeding lifespan for (A) female ($N$ = 120) and (B) male ($N$ =120) breeders, and the effects of distance moved for later pairings, relatedness to mate, and breeding lifespan on a subset of (C) female ($N$ = 50) and (D) male ($N$ = 91) breeders who switched mates at least once in their lifetime.

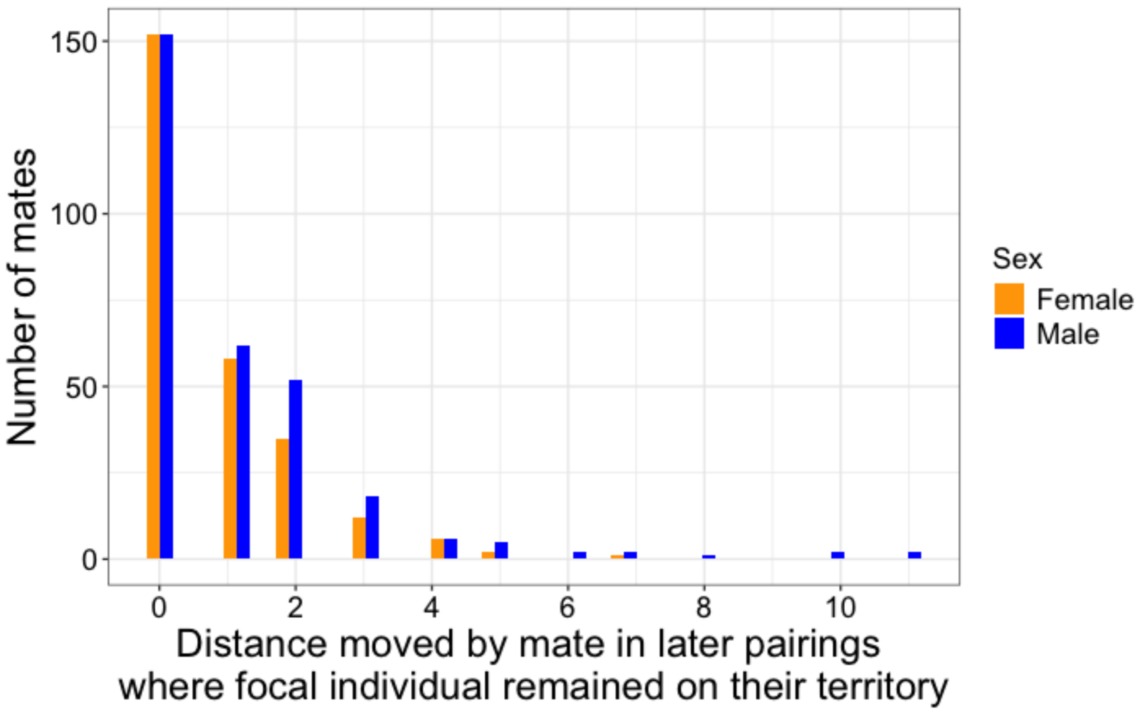


**Figure S6.** Histogram showing distribution of distances moved by new mates of focal individuals who retained their breeding territory for later pairings (orange = female mates, blue = male mates). More than 90% of mates of both sexes dispersed from ≤ 2 territories away.

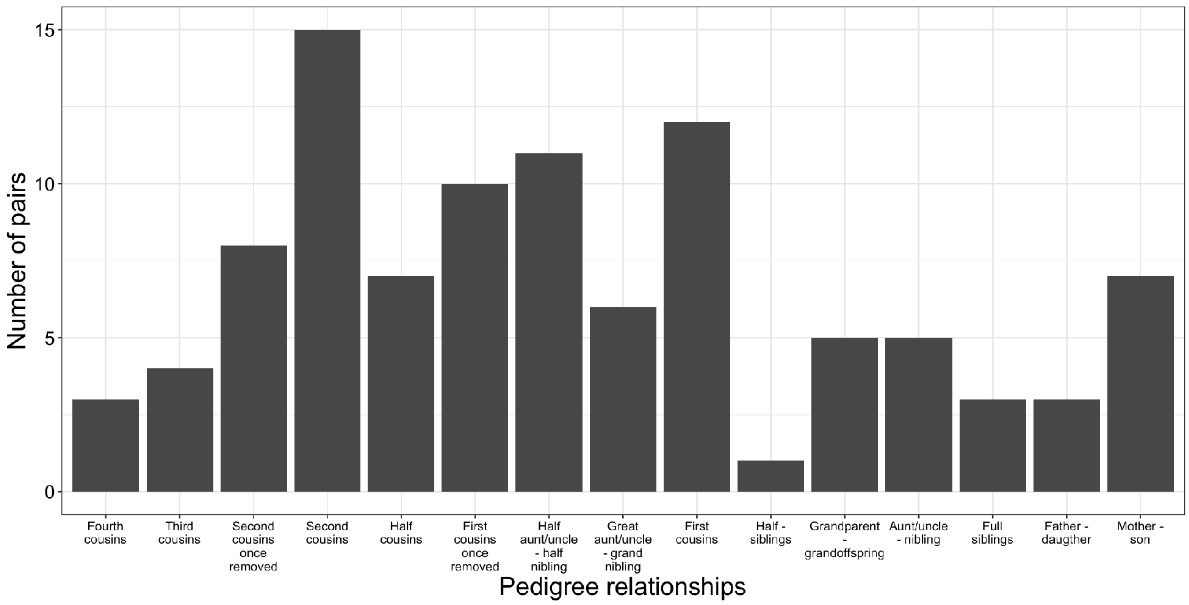


**Figure S7.** The number of Florida scrub-jay breeding pairs consisting of pedigreed relatives in our study population between 1990-2021. Pedigree relationships are arranged in order of increasing relatedness on the x-axis.

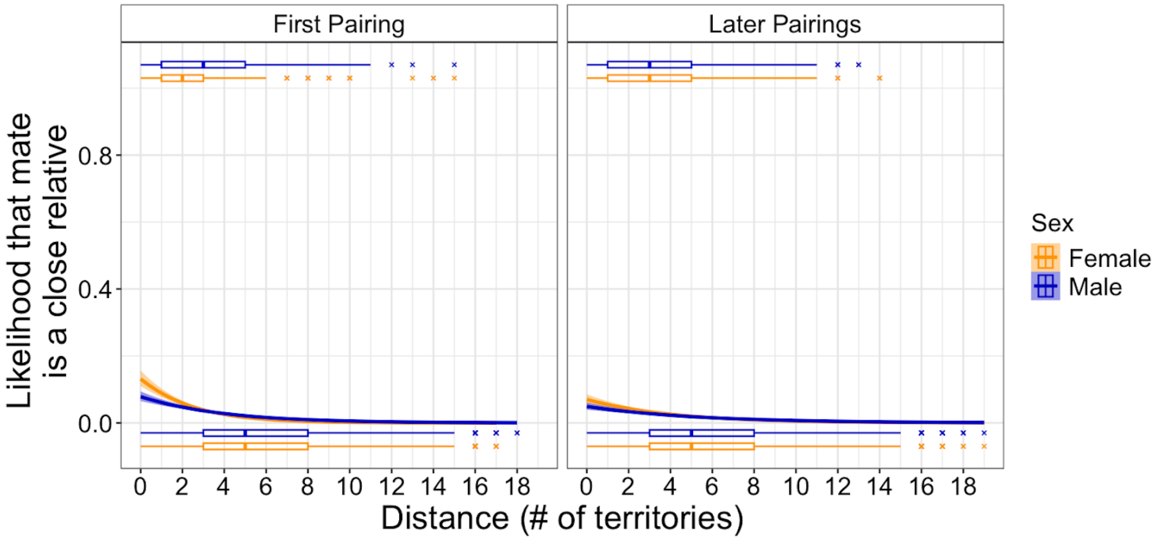


**Figure S8.** For both male (blue) and female (orange) Florida scrub-jays, mates at shorter distances from both their natal ("First Pairings") and breeding territories ("Later Pairings") are more likely to be close relatives. Lines indicate model fits, shaded areas indicate 95% confidence intervals, and boxplots indicate distribution of raw data, with points indicating outliers. We defined close relatives as opposite-sex individuals that are at least as closely related as first cousins to the focal individuals (1 = close relative, 0 = not close relative). We included all instances in which individuals become breeders for the first time that year or switched mates (*N*: Males, First pairing = 12,111; Males, Later pairing = 11,121; Females, First pairing = 10,800; Females, Later pairing = 10,464 possible pairings, with Males, First pairing = 426, Males, Later Pairing = 241, Females, First pairing = 395, and Females, Later pairing = 251 unique individuals).

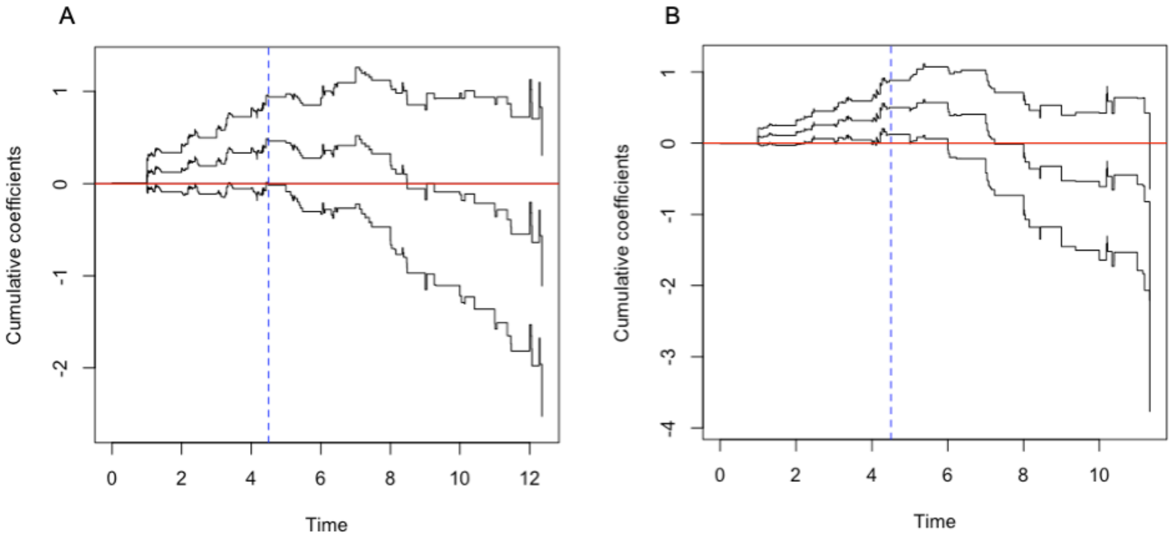


**Figure S9.** Results of time-varying proportional hazards models estimating the effect of distance moved for first pairing on breeder survival for (A) male and (B) female Florida scrub-jays. Between 4 and 5 years post breeding onset, the effect changes from a positive effect on mortality (to the left of the dotted blue line) to a negative effect (to the right of the dotted blue line). The middle black line shows the cumulative coefficient estimate and the top and bottom black lines show the 95% confidence intervals.

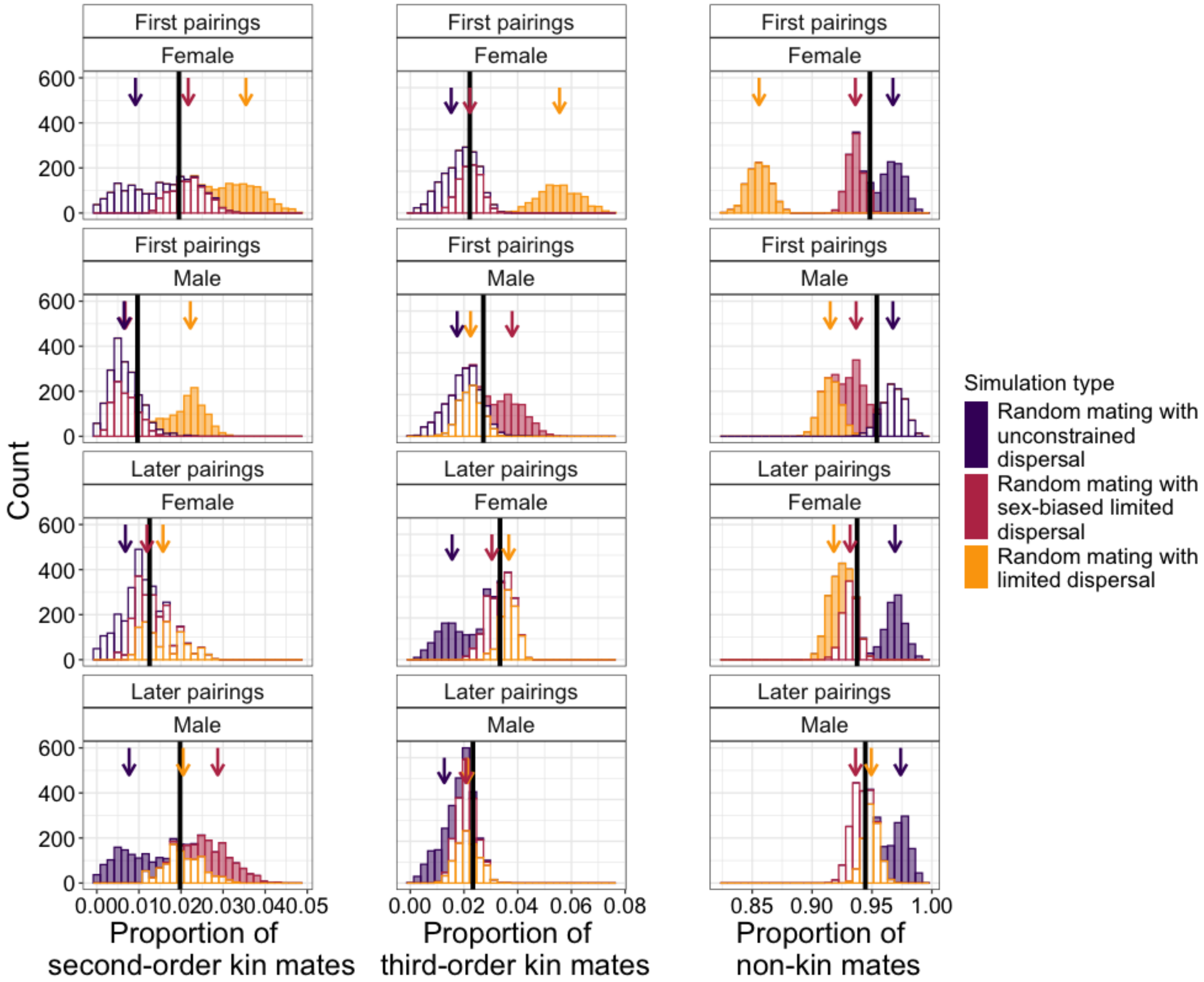


**Figure S10.** Proportion of second-order kin, third-order kin, and non-kin mates observed and expected under three different models of random mating in Florida scrub-jays at two life stages (first and later pairings) and for both sexes (male and female). The observed population means across all 32 years of our study period are indicated by black vertical lines. Histograms show the distributions of expected means under 1000 simulations of random mating in a scenario with unconstrained dispersal (purple), limited and sex-biased dispersal (red), and limited dispersal without sex-bias (yellow) (filled bars = distribution significantly different; unfilled bars = distribution not significantly different from the observed mean). Means of distributions are indicated by arrows.

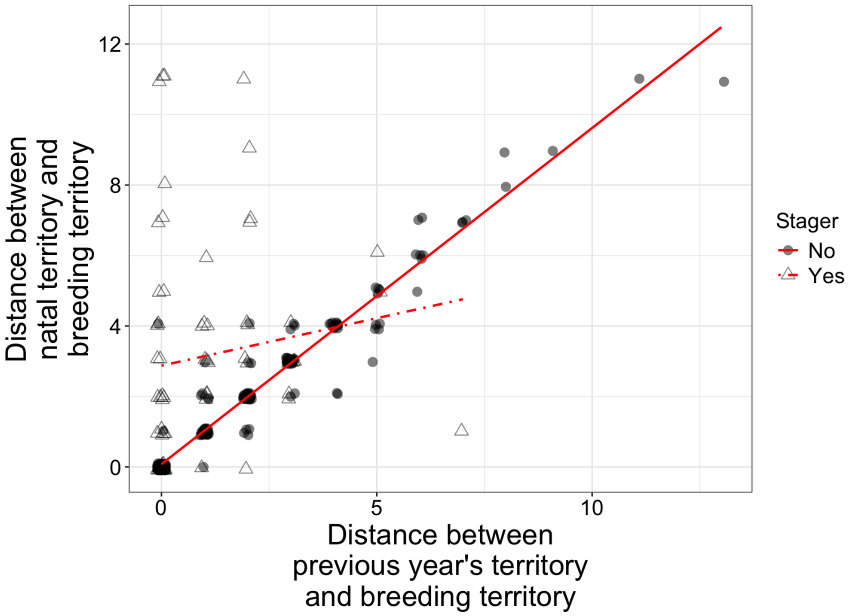


**Figure S11.** The correlation (Pearson's $r$ = 0.70) between the distance between an individual's previous year's territory and breeding territory (x-axis) and the distance between an individual's natal territory and breeding territory (y-axis) for all individuals with lifetime fitness data ($N$ = 317). As expected, the two distances were less correlated for individuals who staged at a non-natal territory before becoming breeders (unfilled triangles; Pearson's $r$ = 0.13, dashed line) than those who remained on their natal territory (filled circles; Pearson's $r$ = 0.97, solid line), though only 21% of individuals staged. Note that territory boundaries shift from year to year, so the number of territories crossed as calculated from the natal territory in the year the individual was born could be different from the number of territories crossed as calculated from that same territory in the year the individual dispersed to become a breeder.

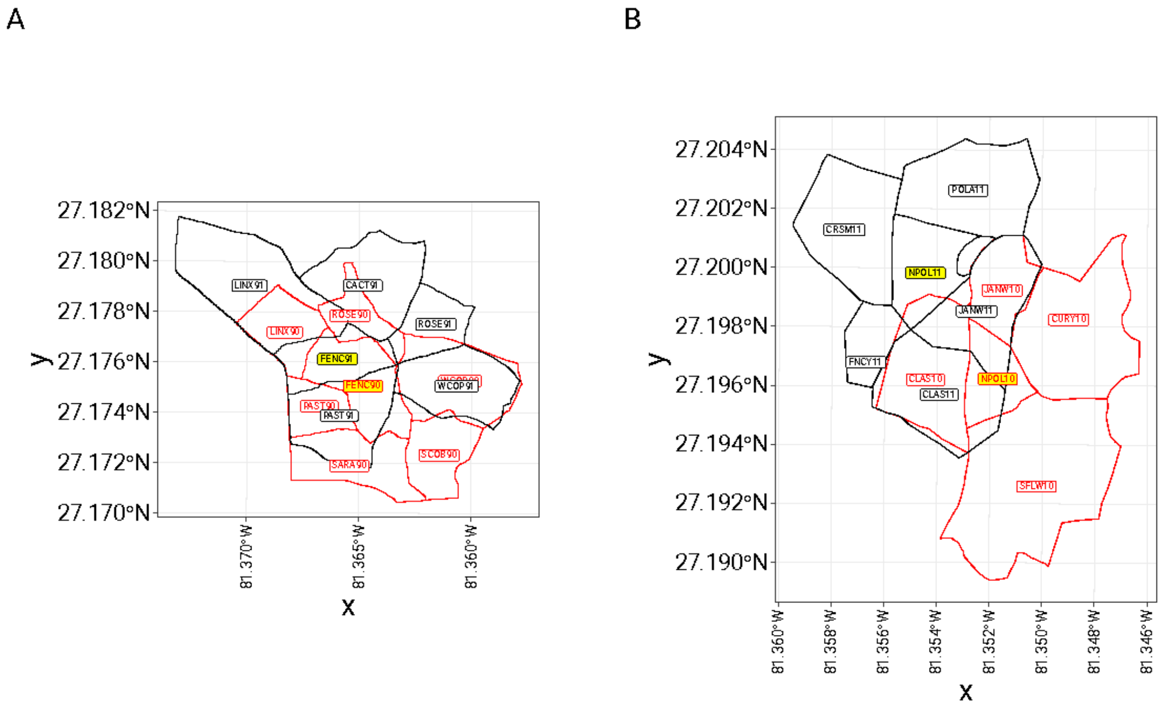


**Figure S12.** When manually calculating distance moved between two territories highlighted in yellow, we assessed the difference between the territory location and configuration between the previous year (red) and the current year (black). (A) When the territory location and configuration with respect to surrounding territories remained largely the same (such as shown here from FENC90 to FENC91), we assigned a distance moved of 0 territories away. (B) When territory location and configuration with respect to surrounding territories changed considerably between the two consecutive years, we calculated distance moved as the number of territories between the previous and current location. For the example shown here (NPOL10 to NPOL11), we assigned a distance moved of 1 territory away.

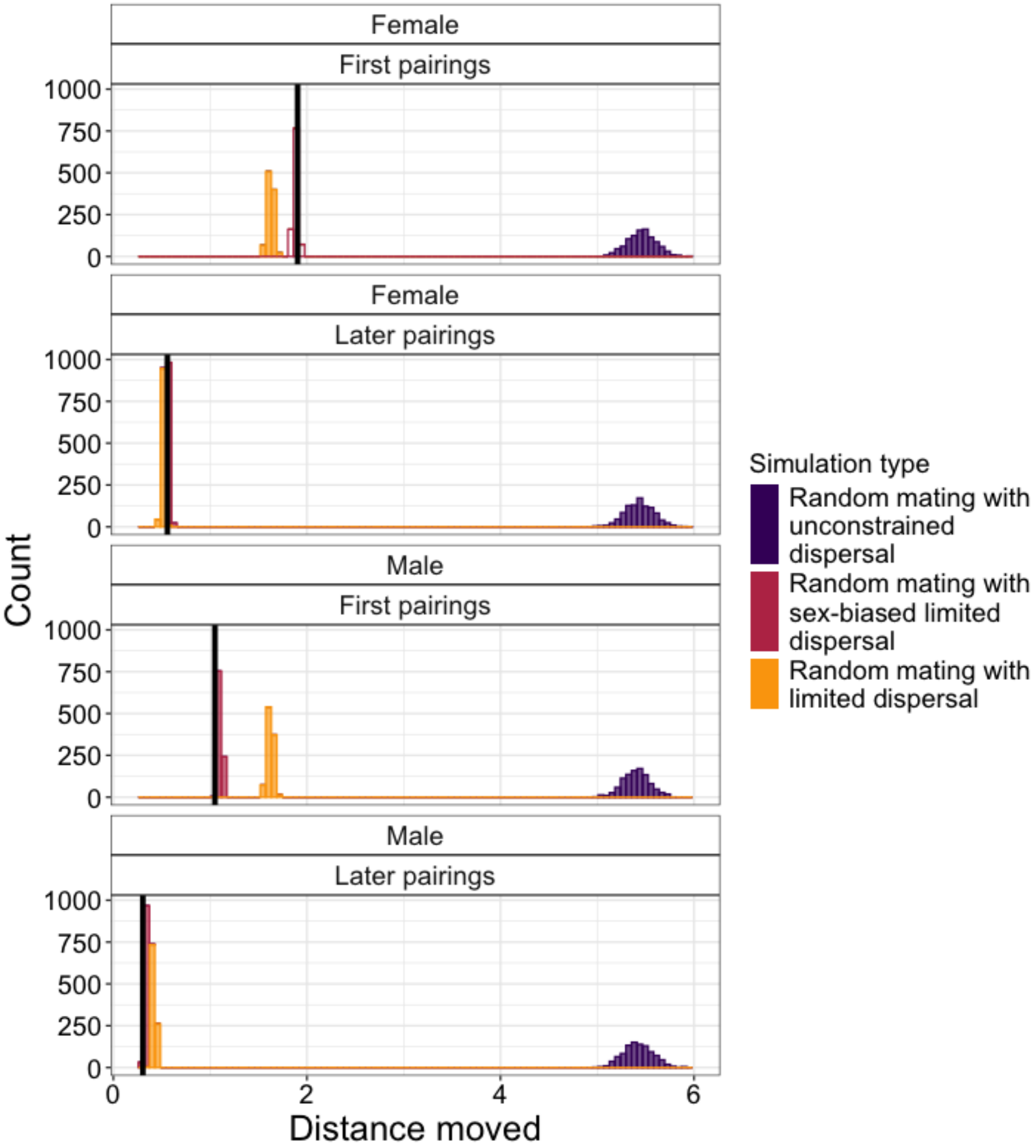


**Figure S13.** Distance moved observed and expected under three different models of random mating in Florida scrub-jays at two life stages (first and later pairings) and for both sexes (male and female). The observed population means across all 32 years of our study period are indicated by black vertical lines. Histograms show the distributions of expected means under 1000 simulations of random mating in a scenario with unconstrained dispersal (purple), limited and sex-biased dispersal (red), and limited without sex-biased dispersal (yellow) (filled bars = distribution significantly different; unfilled bars = distribution not significantly different from the observed mean).

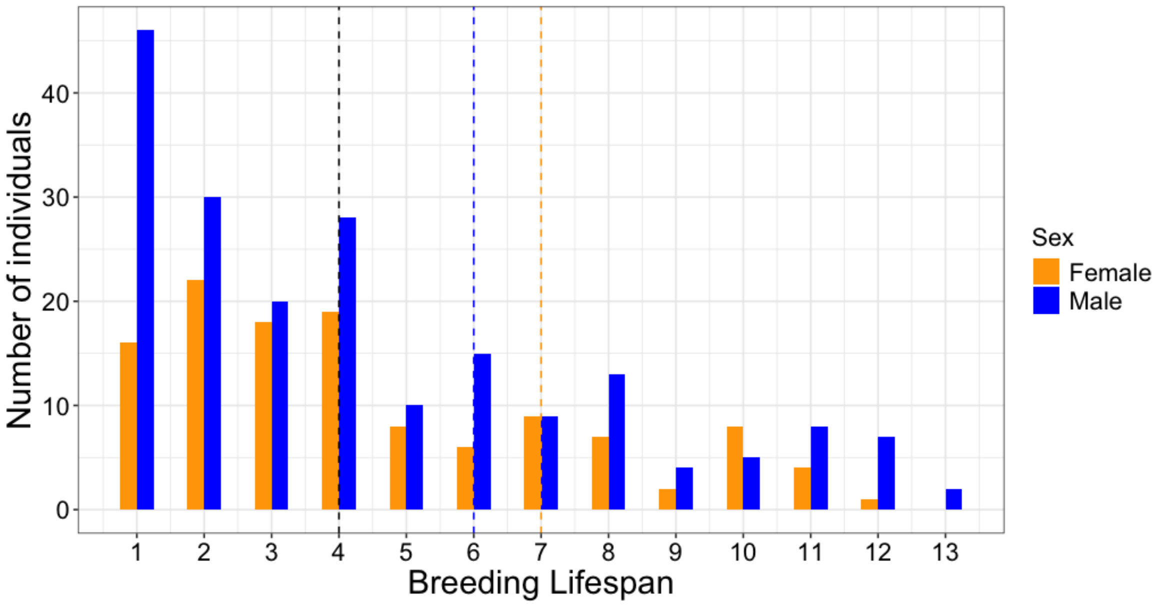


**Figure S14.** The distribution of kinship coefficients for potential mates for both sexes (female = orange, male = blue) of breeding Florida Scrub-Jays in our study population. The dotted vertical line denotes the threshold above which individuals are considered close relatives (0.0625; at least as related as first cousins).

**Table S1.** Results of stratified Cox proportional hazards models estimating the effect of distance moved for first pairing on breeder survival in Florida scrub-jays (*N*: Males = 197, Females = 120), with two time intervals split by cut points at 5, 6, and 7 years post breeding onset. Significant effects are in bold ($P \leq 0.05$).

| **Cut point** | **Sex** | **Fixed Effect** | **Estimate** | **Std Err** | ***Z*** | ***P*** |
|---|---|---|---|---|---|---|
| 5 years | Male | **Intercept** | 0.07 | 0.04 | 1.67 | 0.10 |
| | | Distance moved for first pairing (≤ 5 years) | -0.12 | 0.07 | -1.77 | 0.08 |
| | | **Distance moved for first pairing (> 5 years)** | **0.09** | **0.03** | **3.54** | **<0.001** |
| | Female | **Intercept** | **-0.24** | **0.11** | **-2.25** | **0.02** |
| | | Distance moved for first pairing (≤ 5 years) | 0.08 | 0.05 | 1.69 | 0.09 |
| | | **Distance moved for first pairing (> 5 years)** | **-0.19** | **0.08** | **-2.41** | **0.02** |
| 6 years | Male | **Intercept** | **0.08** | **0.02** | **3.28** | **0.001** |
| | | **Distance moved for first pairing (≤ 6 years)** | **-0.24** | **0.12** | **-2.10** | **0.04** |
| | | Distance moved for first pairing (> 6 years) | 0.07 | 0.05 | 1.46 | 0.15 |
| | Female | **Intercept** | **-0.21** | **0.09** | **-2.28** | **0.02** |
| | | **Distance moved for first pairing (≤ 6 years)** | **0.06** | **0.02** | **2.31** | **0.02** |
| | | Distance moved for first pairing (> 6 years) | -0.18 | 0.15 | -1.22 | 0.22 |
| 7 years | Male | **Intercept** | 0.07 | 0.04 | 1.67 | 0.10 |
| | | Distance moved for first pairing (≤ 7 years) | -0.12 | 0.07 | -1.77 | 0.08 |
| | | **Distance moved for first pairing (> 7 years)** | **0.09** | **0.03** | **3.54** | **<0.001** |
| | Female | **Intercept** | **-0.24** | **0.11** | **-2.25** | **0.02** |
| | | Distance moved for first pairing (≤ 7 years) | 0.08 | 0.05 | 1.69 | 0.09 |
| | | **Distance moved for first pairing (> 7 years)** | **-0.19** | **0.08** | **-2.41** | **0.02** |

**Table S2.** Results of a generalized linear mixed model with a zero-inflated Poisson distribution estimating the effect of total distance moved for later pairings, relatedness to mate, and breeding lifespan on lifetime reproductive success (*i.e.*, number of offspring surviving to adulthood) of Florida scrub-jay male breeders who switched mates at least once in their lifetime (*N*=91). The results were not qualitatively different from a model using a negative binomial error distribution (see *Main Text*). Significant effects are in bold ($P \leq 0.05$).

| **Model component** | **Fixed effect** | **Estimate** | **Std Err** | ***Z*** | ***P*** |
|---|---|---|---|---|---|
| Count model | **Intercept** | 1.40 | 0.07 | 19.24 | **<0.001** |
| | Total distance moved for later pairings | -0.06 | 0.04 | -1.40 | 0.16 |
| | **Weighted kinship coefficient to mates** | -0.41 | 0.10 | -3.93 | **<0.001** |
| | **Breeding Lifespan** | 0.51 | 0.05 | 9.65 | **<0.001** |
| Zero-inflation model | **Intercept** | -2.46 | 0.47 | -5.20 | **<0.001** |
| | Breeding Lifespan | -0.63 | 0.51 | -1.24 | 0.21 |

**Table S3.** Pearson's correlation coefficients between fixed effects included together in models of LRS.

| | Distance moved for first pairing | Weighted kinship coefficient to mates | Breeding lifespan | Total distance moved for later pairings |
|---|---|---|---|---|
| **Distance moved for first pairing** | - | -0.11 | -0.07 | - |
| **Weighted kinship coefficient to mates** | - | - | 0.02 | -0.01 |
| **Breeding lifespan** | - | - | - | - |
| **Total distance moved for later pairings** | - | - | - | - |

**Table S4.** Results of generalized linear mixed models estimating the effect of distance between natal and first breeding territory, relatedness to mate, and breeding lifespan on lifetime reproductive success (*i.e.*, number of offspring surviving to adulthood) of Florida scrub-jay breeders. Significant effects are in bold ($P \leq 0.05$).

| Dataset | Sex | Fixed Effect | Estimate | Std Err | *Z* | *P* |
|---|---|---|---|---|---|---|
| All breeders with lifetime data between 1990-2021 | Male ($N$=197) | **Intercept** | **0.47** | **0.08** | **5.66** | **<0.001** |
| | | Distance between natal territory and first breeding territory | -0.04 | 0.10 | -0.42 | 0.68 |
| | | **Weighted kinship coefficient to mates** | **-0.37** | **0.12** | **-3.20** | **0.002** |
| | | **Breeding Lifespan** | **0.93** | **0.06** | **15.07** | **<0.001** |
| | Female ($N$=120) | **Intercept** | **0.73** | **0.10** | **7.48** | **<0.001** |
| | | Distance between natal territory and first breeding territory | -0.13 | 0.08 | -1.62 | 0.10 |
| | | Weighted kinship coefficient to mates | -0.04 | 0.09 | -0.51 | 0.61 |
| | | **Breeding Lifespan** | **0.85** | **0.09** | **9.44** | **<0.001** |

**Table S5.** Staging duration (in years) inferred using data from Suh et al. 2022. The median duration of staging for both sexes was 2 years.

| **Data from Suh et al. 2022** | | | **Our calculation based on data from Suh et al. 2022** | |
|---|---|---|---|---|
| **Sex** | **Year of life** | **Number of stagers** | **Number of years staging** | **Number of stagers** |
| Male | 1 | 50 | 1 | 50-39 = 11 |
| | 2 | 39 | 2 | 39-12 = 27 |
| | 3 | 12 | 3 | 12-5 = 7 |
| | 4 | 5 | 4 | 5-3 = 2 |
| | 5 | 3 | 5 | 3-1 = 2 |
| | 6 | 1 | 6 | 1-0 = 1 |
| Female | 1 | 50 | 1 | 50-29 = 21 |
| | 2 | 29 | 2 | 29-6 = 23 |
| | 3 | 6 | 3 | 6-3 = 3 |
| | 4 | 3 | 4 | 3-1 = 2 |
| | 5 | 1 | 5 | 1-0 = 1 |
